# CD1a-Mediated Presentation of Canonical Microbial Peptides to T Cells

**DOI:** 10.64898/2026.05.05.723095

**Authors:** Bruno Jorge de Andrade Silva, Annemieke de Jong, Linda A. Fischbacher, Maria Angela M. Marques, Annaliza Legaspi, Adam Shahine, Jade Kollmorgen, Peter A. Sieling, Aaron Choi, Hee Jin Kim, Carlos Adriano Matos e Silva, Kristofor J. Webb, Jason Bradshaw, Patrick J. Brennan, Alina Marusina, Khiem A. Tran, Euzenir Nunes Sarno, Roberta Olmo Pinheiro, Dirk M. Zajonc, D. Branch Moody, Kayvan R. Niazi, Emanual Maverakis, Alessandro Sette, Jamie Rossjohn, Maria T. Ochoa, John T. Belisle, Robert L. Modlin

## Abstract

Langerhans cells express the nonpolymorphic antigen-presenting molecule CD1a, positioning them as contributors to host immunity against *Mycobacterium leprae* in human leprosy. CD1a was originally shown to present non-canonical lipopeptide antigens such as dideoxymycobactin and chemically diverse hydrophobic ligands. Here, we generated CD4⁺ T cell lines from leprosy lesions that recognized *M. leprae* in a CD1a-restricted manner. Unexpectedly, antigen recognition was protease-sensitive, prompting biochemical purification that identified two microbial protein antigens: LppX, a 25-kDa lipoglycoprotein, and Ag85A, a 30-kDa secreted protein with no known lipid modification. Recombinant proteins activated the corresponding T cell lines in a CD1a-dependent manner. Epitope mapping identified 12-mer peptides that fully reconstituted antigenicity, were conserved between *M. leprae* and *M. tuberculosis*, and elicited robust, dose-dependent IFN-γ production and T cell proliferation, establishing that DNA-encoded, ribosomally translated peptides serve as CD1a-restricted cognate antigens. Biochemical analyses showed peptide binding to CD1a, supported by isoelectric focusing and surface plasmon resonance (*K*_D_ ∼75 μM for Ag85A). CD1a–peptide tetramers specifically stained cognate T cells, soluble CD1a was sufficient to present peptide antigen, and transfer of the LppX-specific TCR into naïve T cells restored antigen responsiveness. Using CD1a–peptide tetramers, we identified antigen-specific T cells enriched in patients undergoing reversal reactions compared with patients with lepromatous leprosy and healthy donors. The CD1a-restricted T cell lines secreted IFN-γ and IL-26, cytokines with established antimicrobial activity. Together, these findings demonstrate that CD1a can present canonical microbial peptides as part of a cell-mediated immune response in leprosy, extending the known spectrum of CD1a ligands. Because CD1a is nonpolymorphic and presents antigens to antimicrobial T cells, CD1a–peptide complexes may provide a broadly applicable platform for studying, detecting, and potentially targeting mycobacterial immunity.

## INTRODUCTION

The human CD1 family consists of MHC class I-like molecules that present antigens to T cells. In contrast to the highly polymorphic MHC molecules, CD1 proteins are essentially monomorphic, showing minimal variation between individuals^1^. The group 1 CD1 proteins (CD1a, CD1b, CD1c) are best known for presenting lipid antigens derived from microbes and self-tissues. Landmark studies in the 1990s demonstrated that CD1-restricted T cells can recognize mycobacterial lipid and glycolipid antigens, such as dideoxymycobactin (CD1a)^2,3^, glucose monomycolate (GMM; CD1b)^4^, mycolic acids (CD1b)^5^, mannosylated lipoarabinomannan (manLAM; CD1b)^6^, or mannosyl phosphodolichol (MPD; CD1c)^7^. Co-crystal structures^8,9^ and recognition studies of lipid analogs^4^ demonstrated that CD1-glycolipid complexes formed such that lipid tails are hidden inside CD1b and carbohydrates protrude to the surface for T cell receptor contact, so that co-recognition of CD1-lipid has parallels of co-recognition of MHC-peptide. CD1b and CD1c present a broad array of bacterial lipids (some requiring endosomal loading pathways)^10–13^.

CD1a has been shown to present *M. tuberculosis* dideoxymycobactin, an unusual lipopeptide siderophore containing a short peptide backbone with modified amino acids and attached alkyl chains to T cells^2^. Thus, this antigen fits the prevailing paradigm of CD1 presenting amphipathic molecules with hydrophobic lipid tails^14^. Notably, CD1a can display certain self-lipids (e.g. skin lipids) and allergen-derived lipids to T cells, contributing to inflammatory skin diseases like atopic dermatitis and contact hypersensitivity^15–18^. Crystal structures of CD1a with bound ligands revealed a binding groove with two hydrophobic pockets (A’ and F’) that accommodate alkyl chains^14,19^. These findings reinforced the understanding that CD1a specializes in presenting amphipathic glycolipids and lipopeptide molecules.

Langerhans cells (LCs) are a specialized subset of skin-resident dendritic cells characterized by co-expression of CD1a and langerin (CD207), with an established role in cutaneous host defense. Early studies demonstrated that CD1a⁺ LCs are preferentially enriched at sites of effective immune control of infection. In tuberculoid (T-lep) leprosy lesions, the self-limited form of *M. leprae* infection, CD1a⁺ LCs are present at significantly higher frequencies than in disseminated lepromatous (L-lep) lesions^20–22^. Similarly, in patients undergoing reversal reactions (RR), a cell-mediated immune upgrading from the lepromatous to the tuberculoid pole, CD1a⁺ LC frequencies resemble those observed in T-lep lesions and exceed those in L-lep lesions^23,24^. Together, these observations suggest that CD1a-restricted immune responses correlate with stronger cell-mediated immunity and are positioned at tissue sites consistent with control of *M. leprae* infection. Indeed, prior studies demonstrated that CD1-restricted T cells can have direct antimicrobial activity. For example, CD1-restricted T cells produce granulysin and other cytolytic molecules that kill intracellular mycobacteria, and they release cytokines (e.g. IFN-γ) that activate infected macrophages and LCs to eliminate bacteria^23,25–31^.

Despite these clues, the actual antigen(s) presented by CD1a at the site of disease during *M. leprae* infection remained unknown, aside from evidence that the activity resides in mycobacterial cell lysates^32^. Given that mycobacterial lipoproteins contain both a peptide backbone and lipid modifications, we initially hypothesized that CD1a-restricted T cells might recognize a peptide epitope within a mycobacterial lipoprotein, with the peptide portion anchored in CD1a by its attached lipids, as occurs for CD1a-dideoxymycobactin^14^ and for a synthetic lipopeptide derivative of the HIV NEF protein with CD1c^33^. Conventional, non-acylated peptides generated by standard proteolysis of proteins had not been definitively shown to be presented by CD1 proteins. In humans, protein antigens are generally presented by HLA-encoded class I and class II molecules, whereas CD1 molecules have been predominantly associated with the presentation of lipids, lipopeptides, glycolipids, and other hydrophobic non-peptide antigens. To test these ideas, we set out to identify the molecules recognized by CD1a-restricted T cells from *M. leprae*-exposed individuals. In this study, we therefore examined CD1a-restricted CD4⁺ T cell lines derived from leprosy patient lesions to determine the nature of their target antigens.

## RESULTS

### CD1a-restricted T cell lines from leprosy lesions recognize mycobacterial antigens

We generated T cell lines from leprosy RR patient skin biopsy samples by co-culturing T cells extracted from the skin lesions with *M. leprae*-pulsed antigen-presenting cells (APCs). As antigen-presenting cells, we used CD1a⁺ langerin⁺ Langerhans cell-like dendritic cells (LCDCs) generated from human CD34⁺ progenitor cells, as described previously^31,32,34^. Multiple CD4^+^ T cell lines were obtained that responded to *M. leprae* sonicates in the presence of CD1a^+^ LCDCs. Three CD4^+^ T cell lines, termed LCD4.G, LCD4.C and LCD4.D, were isolated for detailed study. To further characterize the antigen specificity, we tested the T cell lines against a panel of bacteria using CD1a^+^ LCDCs as antigen presenting cells (**Figure 1A**). LCD4.G, LCD4.C and LCD4.D responded strongly to sonicates of *M. leprae* (the leprosy bacillus) and *M. tuberculosis*. LCD4.G exhibited a more restricted reactivity against mycobacterial species than LCD4.C and LCD4.D, with specificity for antigens shared by *M. leprae* and *M. tuberculosis* but absent in the non-pathogen *M. smegmatis* (**Figure 1A**). In contrast, LCD4.C and LCD4.D exhibited comparable levels of response to *M. leprae*, *M. tuberculosis*, and *M. smegmatis*.

**Figure 1.**
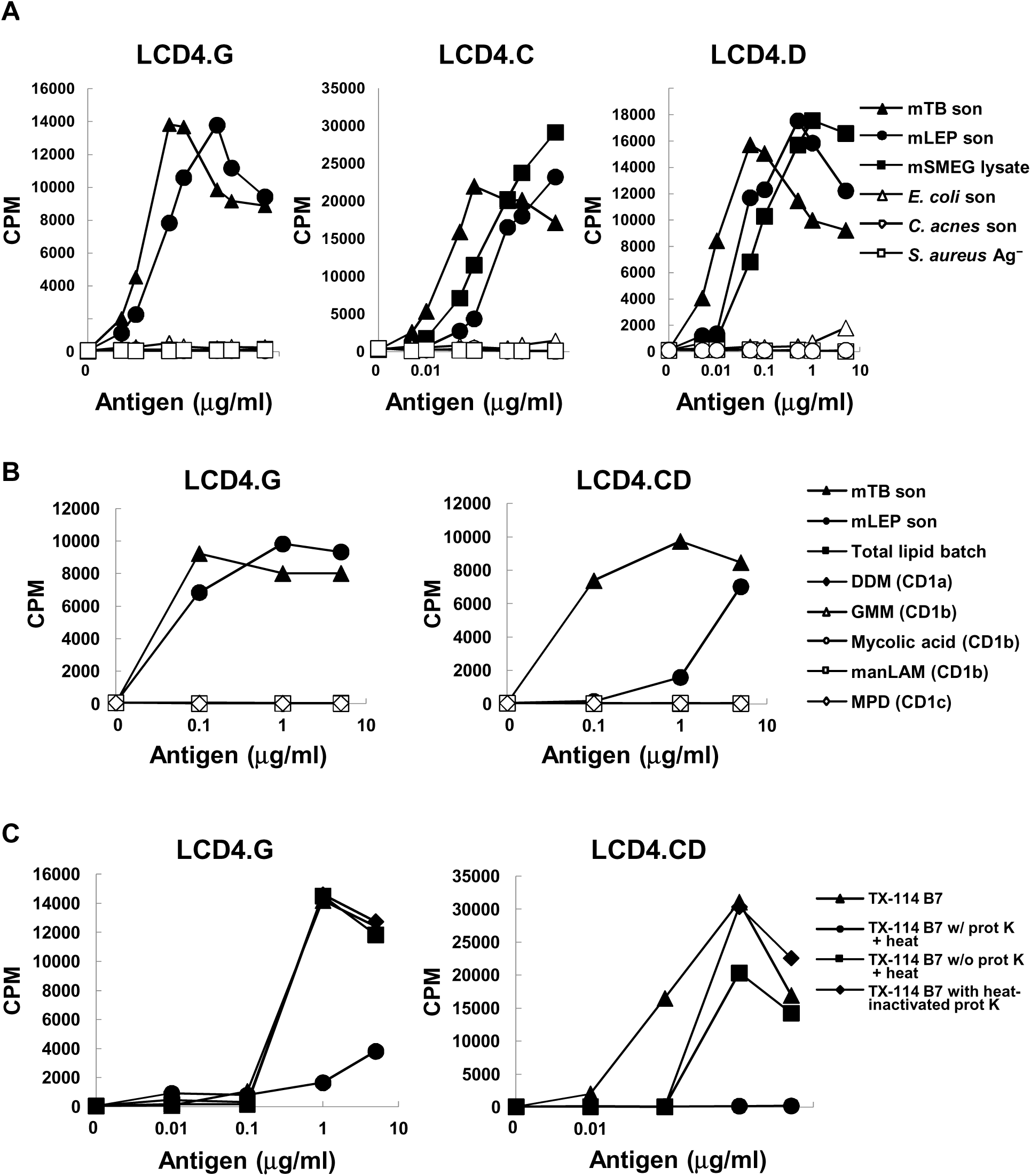
CD1a-restricted T cells recognize protease-sensitive, mycobacteria-specific antigens. (A) LCD4.G, LCD4.C, and LCD4.D T cell lines responses to *M. tuberculosis* (mTB son), *M. leprae* (mLEP son), *M. smegmatis* (mSMEG) and non-mycobacterial lysates. (B) LCD4.G and LCD4.CD T cell lines were tested for recognition of known CD1-presented antigens, including dideoxymycobactin (DDM), glucose monomycolate (GMM), mycolic acid, mannose-capped lipoarabinomannan (manLAM), and mannosyl phosphodolichol (MPD). (C) Effect of proteinase K and heat treatment of the *M. tuberculosis* Triton X-114 extract on T cell proliferation. TX-114 extract was added to the T cell assay after one of four treatments: untreated; proteinase K treatment followed by heat inactivation; heat treatment alone; or incubation with heat-inactivated proteinase K. Data indicate mean ± SEM of triplicate values and are representative of three independent experiments (A–C).

These differences implied that the T cell lines recognize distinct mycobacterial antigens. By comparison, all the T cell lines showed little response to sonicates of unrelated bacteria such as *Escherichia coli*, *Cutibacterium acnes*, or *Staphylococcus aureus* (**Figure 1A**).

To further explore the T cell specificity, TCRα and TCRβ sequences were obtained from the three T cell lines: LCD4.G, LCD4.C, and LCD4.D (**Figure S1A**). LCD4.G expressed a unique TCRα chain composed of TRAV12-2 and TRAJ31 gene segments with a distinct CDR3α sequence: CAVNPPNNARLMF. This sequence differed from the other lines at both the nucleotide and amino acid levels, indicating that LCD4.G expresses a distinct TCR. In contrast, LCD4.C and LCD4.D shared an identical TCRα chain, using TRAV29DV5 and TRAJ41, with the CDR3α sequence: CAASSNSGYALNF. Both the V-J pairing and the CDR3 nucleotide sequences were identical, confirming that these two lines express the same TCR. Analysis of the TCRβ chains revealed a similar pattern. LCD4.G expressed a distinct TRBV11-2/TRBJ1-1 TCRβ chain with the CDR3β motif: CASRPLGGGTNEKLFF. Whereas LCD4.C and LCD4.D shared an identical TRBV16/TRBJ2-1 TCRβ chain encoding: CASSPRRITPYNEQFF. In summary, LCD4.G represented a distinct TCR specificity and clonotype. In contrast, LCD4.C and LCD4.D shared an identical TCRαβ pair and VDJ region, representing a clonal population likely derived from the same T cell precursor, with one clonotype expanded across multiple cells. In subsequent studies, these lines were used interchangeably and are referred to collectively as LCD4.CD.

To determine whether recognition of mycobacterial antigens was CD1a-restricted, both T cell lines were assessed using anti-CD1a blocking antibodies versus an isotype control in co-cultures of T cells, CD1a^+^ LCDCs and sonicated *M. tuberculosis*. Blocking with anti-CD1a antibodies resulted in robust inhibition of proliferation and IFN-γ release in both T cell lines as compared to isotype control (**Figure S1B–S1E**). Overall, for the LCD4.G T cell line, CD1a blockade significantly inhibited the response to *M. tuberculosis* by 61.8% ± 6.5 (p < 0.01) (**Figure S1D**). Similarly, in the LCD4.CD T cell line, CD1a blockade (**Figure S1E**) resulted in percent inhibition of 66.8% ± 12.1 (p < 0.01). In contrast, treatment with an IgG1 isotype control antibody caused minimal inhibition, 16.2% ± 4.6 in LCD4.G and 7.7% ± 3.8 in LCD4.CD, neither of which differed significantly from the *M. tuberculosis* condition. Together, these results demonstrate that both T cell lines are activated by mycobacterial antigens in a CD1a-restricted manner. Given the requirement for an antigen-presenting molecule for T cell activation, we inferred that the mechanism likely involved an antigen binding to CD1a and TCR recognition, rather than direct recognition by TLR or similar mechanisms, which guided later experiments to test the mechanism.

The T cell lines did not recognize previously described CD1-presented mycobacterial antigens (**Figure 1B**). Neither line responded to dideoxymycobactin (DDM; CD1a), glucose monomycolate (GMM; CD1b), mycolic acids (CD1b), mannosylated lipoarabinomannan (manLAM; CD1b), or mannosyl phosphodolichol (MPD; CD1c). Additionally, a total lipid extract of *M. tuberculosis* failed to activate the T cell lines. These data indicate that the antigens recognized by these cells are distinct from known CD1 ligands.

### CD1a-restricted T cell lines from leprosy lesions recognize peptide antigens

To identify the target antigens recognized by LCD4.G and LCD4.CD, we performed biochemical fractionation and proteomic approach. Because *M. leprae* bacilli cannot be cultured in vitro, we leveraged *M. tuberculosis* as a source of cross-reactive antigen. A detergent extract of *M. tuberculosis* (using Triton X-114) was prepared to enrich for amphipathic molecules such as lipoproteins, lipopeptides and lipoglycans^35,36^. The antigenic activity of this *M. tuberculosis* Triton X-114 extract was destroyed by proteinase K treatment, but was not heat labile, indicating protease sensitivity (**Figure 1C**). The protease sensitivity further distinguished the antigen from previously known CD1a-presented lipids, which are typically protease-resistant but sensitive to lipid extraction^2,5–7^. These initial findings suggested that the CD1a-restricted antigen recognized by the T cell lines is a protein, peptide or lipopeptide rather than a glycolipid. We next investigated whether the LCD4.G and LCD4.CD T cell lines respond to different subcellular fractions of *M. leprae* (**Figure S2A**). Robust T cell proliferation was observed in response to the cytosolic (MLSA), membrane (MLMA), and cell wall (MLCwA) fractions, as well as to the *M. leprae* sonicated (mLEP son) fraction. In contrast, both T cell lines responded only weakly to stimulation with the mycolyl-arabinogalactan-peptidoglycan (mAGP) cell wall core, indicating that the dominant antigens recognized by these CD4⁺ T cells are protein rich subcellular fractions.

Proteomics approaches to identify T cell antigens^37,38^ were applied to identify the specific CD1a restricted antigens. Specifically, the *M. tuberculosis* Triton X-114 extract was resolved by preparative isoelectric focusing (IEF), and resulting fractions assayed for T cell reactivity, and adjacent reactive fractions combined into a single pool. The pool was separated by SDS-PAGE, and a T cell “Western blot” assay was performed to locate T cell stimulatory activity (**Figure S2B**)^37^. LCD4.G and line LCD4.CD each recognized distinct molecular mass regions of the IEF fraction, with minimal overlap, indicating that they target different antigenic molecules (**Figure 2A**).

**Figure 2.**
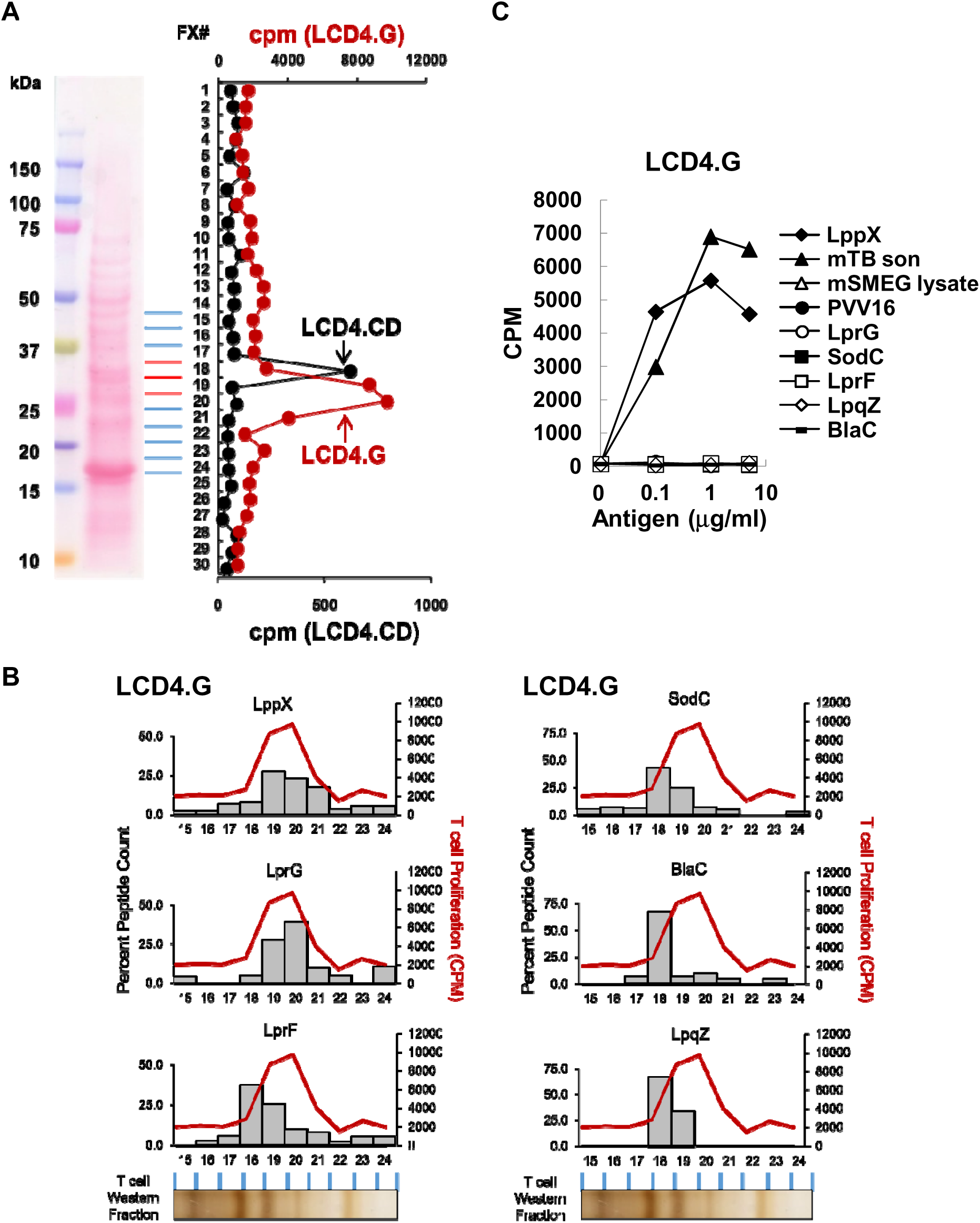
Identification of LppX and Ag85 proteins as CD1a-restricted antigen. (A) The *M. tuberculosis* Triton X-114 extract was subjected to preparative IEF, and each fraction was analyzed by Western blot and screened for T cell activation. (B) Identification of candidate proteins by proteomic analysis (LC–MS/MS) of the active fractions. Active fractions stimulating each T cell line were identified by T cell “Western blot” assay. (C) Proliferative response of LCD4.G to purified recombinant lipoproteins derived from the 25-kDa *M. tuberculosis* lipoglycoprotein LppX. mTB son, *M. tuberculosis* sonicated. mSMEG, *M. smegmatis*. Data indicate mean of triplicate values for LCD4.G T cell line and are representative of three independent experiments (A–C). See also Figures S2 and S5.

LC–MS/MS based proteomics of replicate T cell “Western blot” fractions that activated LCD4.G and adjacent fractions was performed. This identified multiple candidate protein antigens in the 20-30 kDa range, six of which were known or putative lipoproteins with *M. leprae* homologues. Multiple peptides derived from the 25-kDa *M. tuberculosis* lipoprotein LppX were highly enriched in the LCD4.G-reactive fractions and showed the strongest correlation with the T cell “Western blot” signal (**Figure 2B**). LppX is a well-characterized lipoglycoprotein present in the mycobacterial cell envelope^39^. To validate the proteomics results, we measured the proliferative response of LCD4.G to all six potential lipoprotein antigens (**Figure 2C**). As a positive control, *M. tuberculosis* sonicate induced proliferation in contrast to WT *M. smegmatis* lysate (negative control). The lack of proliferation to the negative control is consistent with the absence of a LppX homologue encoded in the *M. smegmatis* genome. In contrast, robust, dose-dependent proliferation was observed in response to recombinant *M. tuberculosis* LppX produced in *M. smegmatis*. Other *M. tuberculosis* lipoproteins, recombinant LprG, SodC, LprF, LpqZ and BlaC failed to activate the LCD4.G T cell line. These results suggest that LppX is the antigen for LCD4.G.

LppX possesses a Sec-translocation signal sequence and the mature protein is acylated and O-glycosylated upon translocation^39,40^. To determine if acylation and/or glycosylation of LppX is required for the response of LCD4.G, we tested recombinant proteins produced in *M. smegmatis* and *E. coli* with and without the signal sequence and relevant bacterial mutants (**Figure S3A**). Again, the WT *M. smegmatis* lysate served as a negative control, the acylated and glycosylated recombinant LppX produced in *M. smegmatis* was the positive control.

Unexpectedly, all recombinant forms of LppX, including the non-acylated forms retained the ability to stimulate LCD4.G. Mass spectrometric analysis of the purified rLppX confirmed that the signal sequence containing form produced in *M. smegmatis* was fully acylated and glycosylated (**Figure S3B**), while the form lacking a signal sequence and produced in *E. coli* was not post-translationally modified (**Figure S3C**). Thus, in contrast to classical CD1a antigens, a lipid tail is not required for the presentation of the LppX epitope and LCD4.G recognizes an unmodified peptide presented by CD1a. Alternatively, because LppX is a lipid carrier protein, LCD4.G might have recognized LppX-associated phthiocerol dimycocerosate rather than the LppX polypeptide itself^39^. However, total lipid extracts of *M. tuberculosis*, which would be expected to contain this lipid family, did not stimulate LCD4.G, and recombinant LppX produced in *E. coli* remained antigenic despite the absence of mycobacterial lipids.

To demonstrate that the epitope presented to LCD4.G was an unmodified peptide, we mapped the origin of the LppX epitope within the polypeptide sequence. Specifically, we generated three overlapping recombinant protein fragments, representing truncated constructs of LppX fused to GST, and tested for their ability to stimulate LCD4.G (**Figure 3A, 3B, and S4A**). Only the LppX-E construct stimulated a T cell response (**Figure 3B**). This smaller protein fragment represented the C-terminus of LppX, further pointing to the direct recognition of the LppX protein sequence as the LCD4.G epitope. We then synthesized overlapping 20-mer peptides covering the C-terminus of LppX (spanning amino acids 147–207 of mature protein, a region unique to LppX). T cell assays revealed that only peptides from the C-terminus were active. Two 20-mer peptides corresponding to residues 172191 and 177196 of mature LppX stimulated LCD4.G, including a 15-mer spanning the common sequence between the two 20-mer peptides (**Figure 3C and 3D**). We designate this peptide spanning residues 177–191, the “LppX_15mer”. Peptides upstream and downstream of these sequences did not. Further truncation of this 20-mer identified the minimal epitope: a 12–amino acid peptide with sequence SHHLVRASIDLG (positions 178–189 of mature LppX) (**Figure 3E and S4B)**. The identification of a synthetic peptide epitope demonstrated the T cell response is directed against peptide rather than LppX-associated lipid. Alignment of the LppX-derived 12-mer peptide from *M. tuberculosis* with the corresponding sequence in *M. leprae* LppX demonstrated complete sequence identity (**Figure 3F; Table S1**), consistent with the ability of LCD4.G to recognize both species.

**Figure 3.**
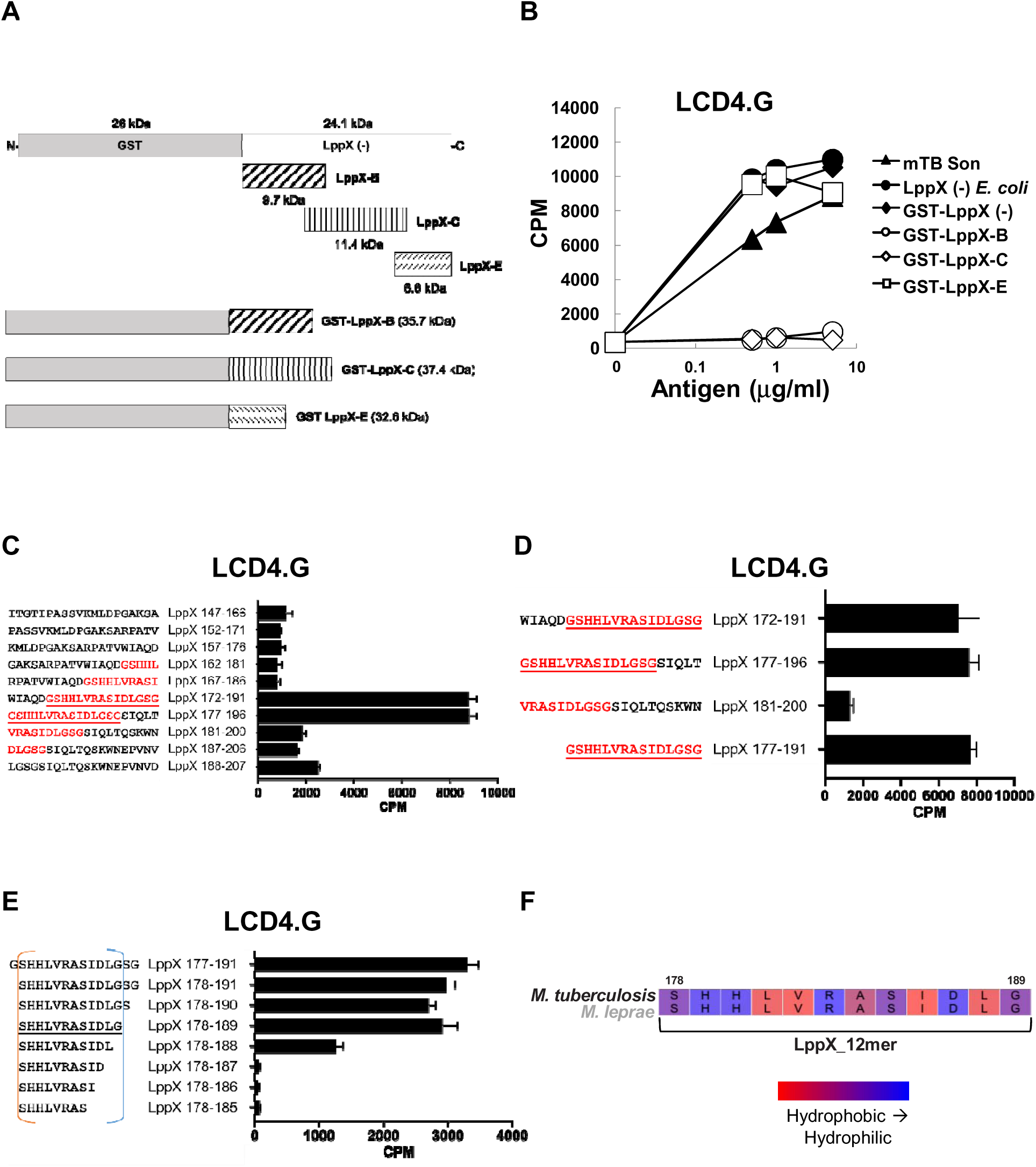
Fine mapping defines the minimal LppX peptide epitope recognized by T cells. (A) Schematic representation of truncated LppX constructs fused to GST. (B) Recombinant GST-LppX fragments spanning the N- and C-termini were evaluated for their capability to stimulate the LCD4.G T cell line. mTB son, *M. tuberculosis* sonicated. Data indicate mean of triplicate values for LCD4.G T cell line and are representative of three independent experiments. (C–D) 20-mer and 15-mer LppX peptide series spanning aa 147–207 were screened for T cell activation. C-terminal truncations of LppX identified 172–191 and 177–196 as active and defined a 15-mer intermediate (aa 177–191). The amino acids corresponding to the LppX_15mer sequence are indicated in red, and sequences containing the full 15-mer are underlined. (E) Overlapping peptides derived from the C-terminus of LppX (15–8 amino acids in length) were screened for their ability to stimulate the LCD4.G T cell line. Truncation analysis identified the minimal 12-aa sequence SHHLVRASIDLG (LppX_12mer) as necessary for T cell activation. The minimal active epitope is underlined. Data indicate mean ± SEM of triplicate values and are representative of three independent experiments (C–E). (F) Alignment of the LppX-derived 12-mer peptide (SHHLVRASIDLG) from *M. tuberculosis* against the corresponding sequence in *M. leprae* LppX. See also Figure S4.

Recognition of a non-acylated peptide epitope by LCD4.G broadened the search criteria applied to T cell Western proteomic data, facilitating identification of the antigen recognized by LCD4.CD. In this case, peptides corresponding to the *M. tuberculosis* Antigen 85 (Ag85) complex were the most abundant in T-cell Western fractions that activated LCD4.CD (**Figure S5A**). Ag85 refers to a family of secreted mycobacterial proteins (∼30 kDa) known for their fibronectin-binding and mycolyl-transferase activity^41^; it includes Ag85A, Ag85B, and Ag85C (also known as FbpA, FbpB, and FbpC respectively). These secreted proteins are neither lipid-modified nor glycosylated. The reactivity was validated using recombinant proteins expressed in *E. coli*, demonstrating that LCD4.CD recognized recombinant *M. tuberculosis* Ag85A and Ag85B, and to a lesser extent, Ag85C (**Figure S5B**). To further define the antigen recognized by LCD4.CD T cells, we next applied the GST-fusion fragment approach to Ag85A, which localized LCD4.CD reactivity to the N-terminal portion of Ag85A (**Figure 4A, 4B, S5C, and S5D**).

**Figure 4.**
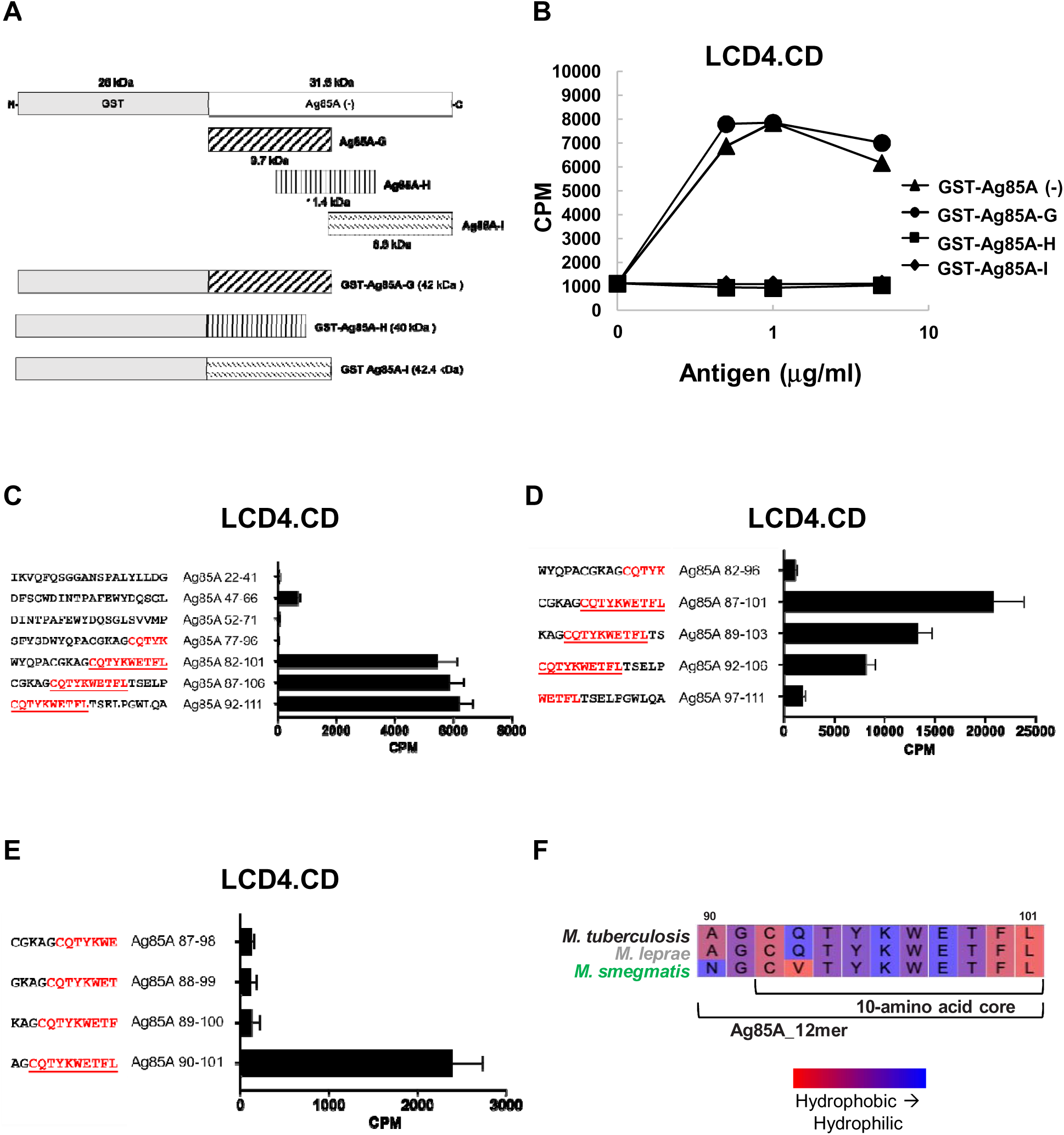
Definition of the minimal Ag85A-derived peptide epitope recognized by T cells. (A) Schematic representation of truncated Ag85A constructs fused to GST. (B) Recombinant GST-Ag85A fragments spanning the N- and C-termini were evaluated for their capability to stimulate the LCD4.CD T cell line. Data indicate mean of triplicate values for LCD4.CD T cell line and are representative of three independent experiments. (C) Selected Ag85A-derived 20-mer candidate peptides from the N-terminal fragment G were tested for activation of LCD4.CD T cells. Reactive peptides localized activity to a CQTYKWETFL-containing region. (D) Overlapping 15-mer peptides spanning the reactive region were tested to refine the epitope, identifying Ag85A residues 87–101 as an active 15-mer. (E) Truncated 12-mer peptides further defined AGCQTYKWETFL as the minimal active 12-mer. The amino acids corresponding to the Ag85A 10-mer shared sequence are indicated in red, and the full shared sequence is underlined (C–E). Data indicate mean ± SEM of triplicate values and are representative of three independent experiments (C–E). (F) Sequence comparison of the Ag85A-derived 12-mer epitope across mycobacterial species. The Ag85A-derived 12-mer peptide from *M. tuberculosis* was aligned with the orthologous Ag85A sequence from *M. leprae* and an Ag85A homolog from *M. smegmatis* (#2078). See also Figure S5.

After localizing LCD4.CD reactivity to the N-terminal fragment of Ag85A, we selected a limited panel of 20-mer candidate peptides for epitope mapping. Candidate selection was informed by comparison of Ag85A and Ag85C amino acid sequences (**Figure S6A**), including regions of sequence divergence, and by qualitative inspection of amino acid composition, guided by features observed in the LppX epitope. Several regions of the fragment were sampled; in particular, a distal region containing clustered sequence differences and mixed residue composition was sampled more densely using overlapping 20-mer peptides. Testing of this candidate panel revealed that only peptides from the distal region stimulated LCD4.CD, whereas peptides from other regions were non-reactive (**Figure 4C**). Alignment of the reactive peptides identified a shared sequence, thereby localizing the epitope to this region.

To further define the epitope, we generated overlapping 15-mer peptides spanning the reactive region and identified a single peptide that retained activity (**Figure 4D**). This 15-mer, corresponding to residues 87–101 of mature Ag85A, was designated the “Ag85A_15mer.” Peptides upstream and downstream showed diminished activity, indicating that recognition was restricted to this region. Further truncation identified a minimal stimulatory 12-mer peptide, AGCQTYKWETFL (residues 90–101), designated the “Ag85A_12mer” (**Figure 4E**). This 12-mer is completely conserved between Ag85A and Ag85B of *M. tuberculosis* but differs in the first four amino acids in Ag85C (**Figure S6B**). Analysis of the overlapping peptides defined a 10–amino acid sequence, CQTYKWETFL, shared among active peptides (**Figure 4C–4E**). This 10-mer is conserved between Ag85A and Ag85B but differs from Ag85C by two amino acids (**Figure S6B**).

Alignment of the Ag85A-derived peptide across mycobacterial species revealed that the CD1a-restricted epitope “Ag85A_12mer” is identical between *M. tuberculosis* and *M. leprae* (**Figure 4F; Table S2**). Since the whole cell lysate of *M. smegmatis* also stimulated LCD4.CD activity (**Figure 1A**), homologous Ag85 sequences in *M. smegmatis* were identified by BLAST analysis using *M. tuberculosis* Ag85A as the query. The *M. smegmatis* homolog MSMEG_2078 was selected based on curated annotation and preservation of residues within the epitope region. The “Ag85A_12mer” of MSMEG_2078 differed by two amino acids within the N-terminus of this 12-mer sequence (A→N and Q→V), while the remaining residues, including the TYKWETFL C-terminal segment, were conserved (**Figure 4F**). Alignment with the homologous region of MSMEG_2078 suggests that limited variation in the N-terminal portion of the Ag85A_12mer is tolerated, whereas the conserved C-terminal TYKWETFL segment is likely critical for recognition. Defining the relative contributions of individual residues to CD1a binding and TCR engagement will require targeted mutational analyses. Further studies are required to determine how the conserved sequence contributes to CD1a presentation and T cell recognition.

These data provide evidence that short peptides derived from microbial proteins can be presented by CD1a to T cells. Notably, the peptide epitopes identified here lack any covalently attached lipid moieties. Together, these findings identify two CD1a-presented microbial peptide antigens, LppX and Ag85A, both derived from mycobacteria and recognized by distinct T cell lines isolated from a leprosy lesion. This challenges the conventional view of CD1 molecules as presenting only lipid antigens, revealing a previously unrecognized role for CD1a in the presentation of peptide antigens to T cells.

### Binding of microbial peptides to CD1a molecules

Having defined the peptide epitopes, we next assessed whether they exhibited physicochemical features compatible with CD1a binding. Because CD1a typically accommodates hydrophobic ligands within its binding groove, we analyzed the hydrophobicity profiles of the LppX_12mer and Ag85A_12mer peptides using the Kyte–Doolittle scale (**Figure S6C**). Both peptides exhibited an amphipathic organization, with interspersed polar and charged residues alongside hydrophobic residues within a constrained 12-mer length. Notably, both peptides terminate in hydrophobic residues (LppX: …LG; Ag85A: …FL) while containing side chains that could remain solvent-exposed, including an aromatic cluster in Ag85A (YKW) and basic and polar residues in LppX (HHR…RS). These features are compatible with models in which hydrophobic residues contribute to CD1a association while polar and charged residues are available for TCR interaction. Although the sequences differ, this shared organization suggests that both peptides are capable of associating with CD1a. The presence of histidine residues in the LppX-derived peptide raises the possibility that pH-dependent protonation could influence peptide behavior, although this was not directly tested.

To assess peptide binding, soluble CD1a (sCD1a), consisting of the extracellular domain of the human CD1a heavy chain heterodimerized with β2-microglobulin, was expressed and purified. sCD1a was incubated with either LppX_12mer or Ag85A_12mer and analyzed by analytical isoelectric focusing (IEF) gel electrophoresis (**Figure 5A**). Both peptides induced distinct shifts in the isoelectric point (pI) of sCD1a relative to the unloaded protein, consistent with peptide association. The IEF assay provided the first biochemical evidence that conventional microbial peptides associate with CD1a molecules.

**Figure 5.**
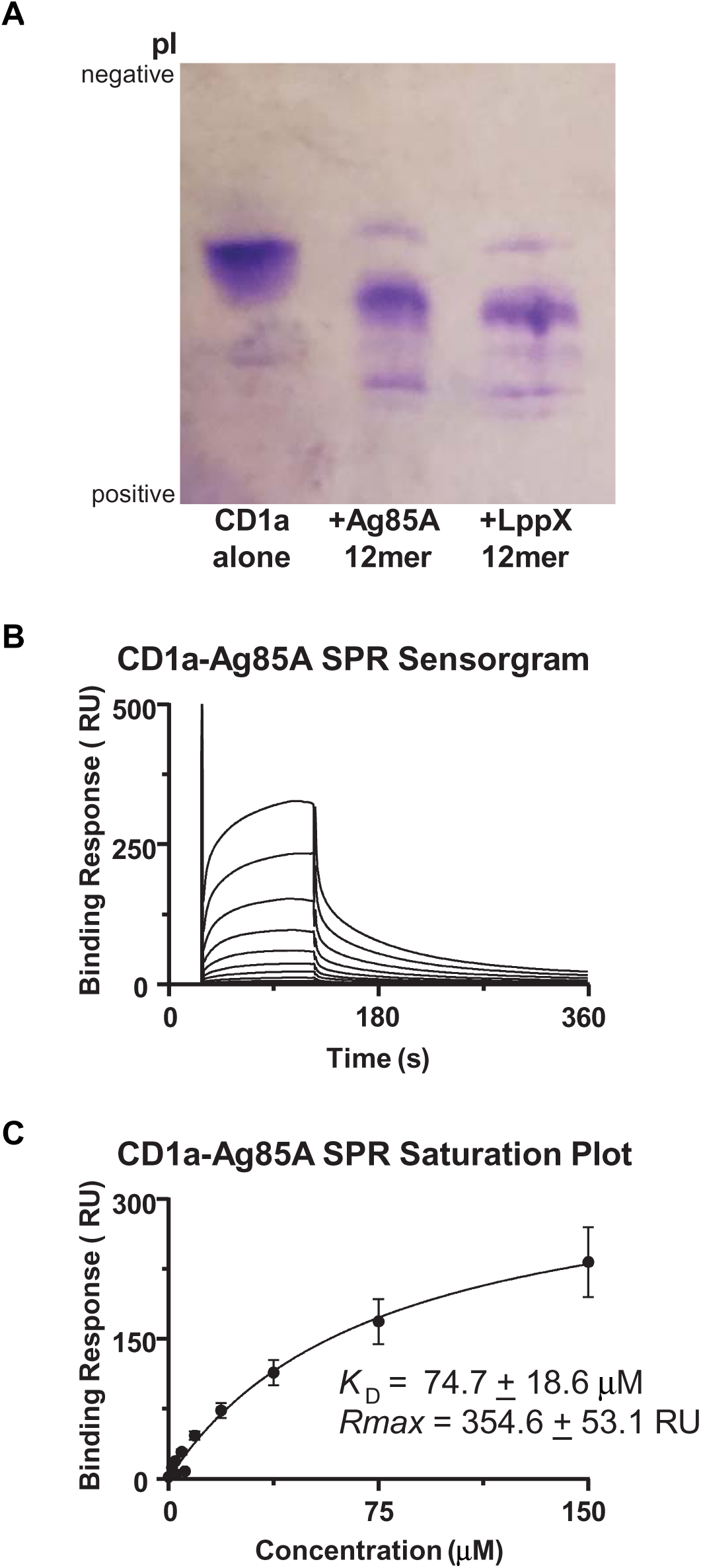
Biochemical and biophysical evidence of peptide–CD1a binding. (A) Analytical isoelectric focusing (IEF) gel of soluble CD1a (sCD1a) alone or following incubation with Ag85A_12mer or LppX_12mer peptides. Position of negative and positive pI charges are indicated. (B–C) Quantitative analysis of Ag85A_12mer peptide binding to sCD1a by surface plasmon resonance (SPR). Representative sensorgram (B) and equilibrium fit (saturation plot) (C). Error bars indicate mean ± SEM from three independent experiments. Dissociation constants (apparent *K*_D_) and Rmax values indicate binding affinity and maximum binding response of CD1a to Ag85A_12mer. See also Figure S6.

### Quantitative measurement of peptide–CD1a binding by SPR

To quantify binding affinities, we performed surface plasmon resonance (SPR) experiments. A biotinylated Ag85A_12mer peptide immobilized on a streptavidin (SA) chip bound sCD1a in a dose-dependent manner, yielding an equilibrium dissociation constant (*K*_D_) of 74.7 μM (**Figure 5B and 5C**), an apparent affinity compatible with biologically relevant antigen-presentation interactions. Because the peptide was immobilized and soluble CD1a was flowed over the surface, this value should be interpreted as an apparent affinity under the assay conditions. CD1a also bound to an LppX_12mer-coated surface, albeit with a lower response (**Figure S6D and S6E**). SPR experiments with the LppX peptide indicated qualitative binding of CD1a at high concentrations, although a reliable *K*_D_ could not be determined. A biotinylated irrelevant control peptide, DQ2.5-glia-α1a, did not bind CD1a (**Figure S6F**). Together with the electrophoretic shift results, these data provide biochemical and biophysical evidence that both microbial peptides bind to CD1a.

### Functional recognition of peptide–CD1a complexes by cognate T cells

Transfer of the LCD4.G TCR into CD4-expressing 5KC reporter cells conferred responsiveness to LppX_12mer presented by CD1a, demonstrating that this TCR is sufficient to mediate LppX–CD1a recognition. 5KC murine T cell hybridomas, which lack endogenous TCR α- and β-chains, were engineered to co-express human CD4 and the LCD4.G TCR (**Figure S6G**). These transfectants produced IL-2 in response to LppX_12mer or LppX produced in *E. coli* (LppX(-)) presented by CD1a⁺ APCs, mirroring IFN-γ production by the parental LCD4.G T cells (**Figure 6A and 6B**). Together, these findings demonstrate that the LCD4.G TCR is sufficient to confer peptide–CD1a recognition.

**Figure 6.**
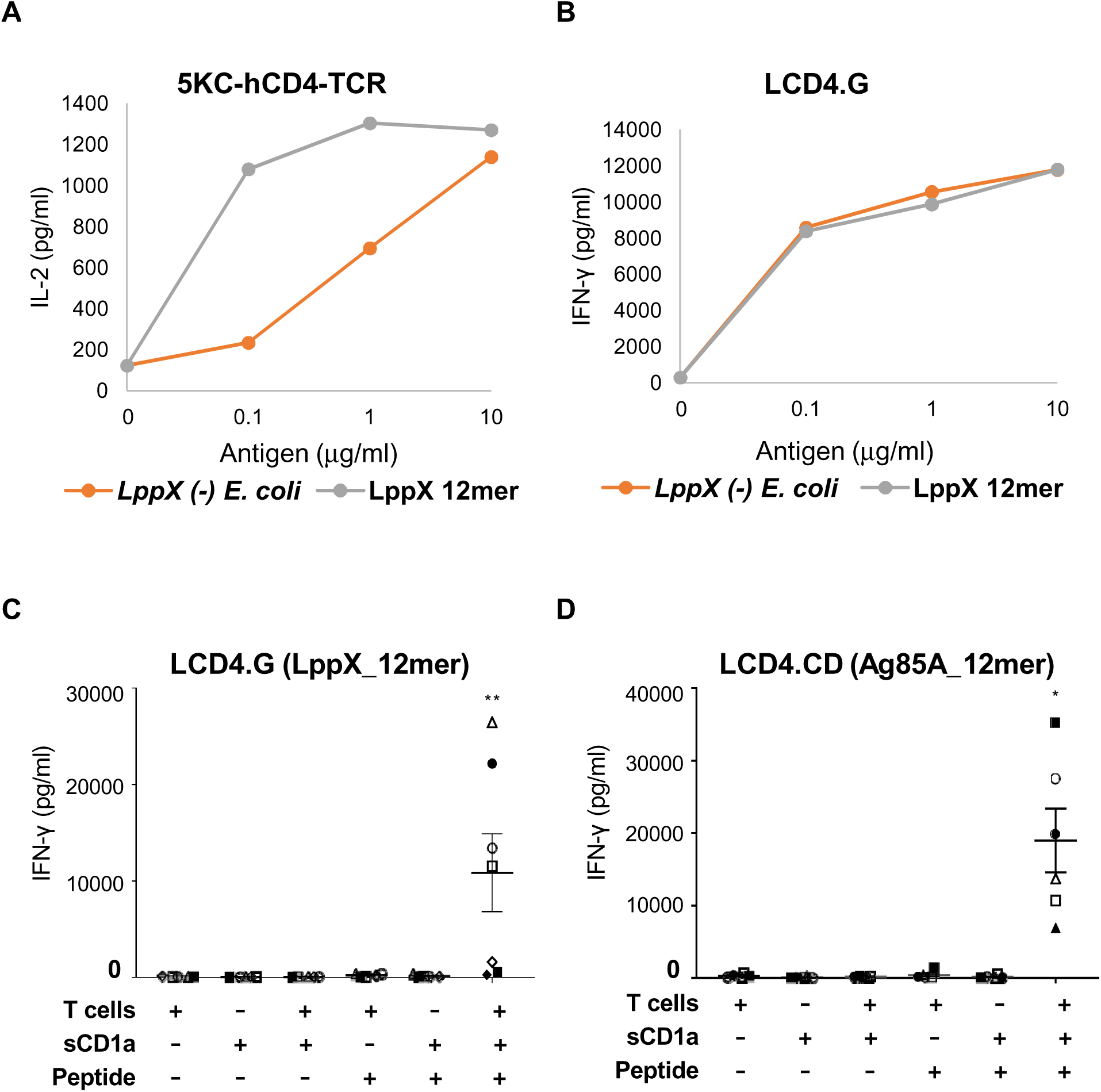
TCR-mediated recognition of peptide–CD1a complexes supports T cell activation. (A) IL-2 secretion by murine 5KC T cell hybridomas expressing human CD4 and the LCD4.G TCR upon stimulation with LppX_12mer or LppX expressed in *E. coli*. (B) IFN-γ release by parental LCD4.G T cells. Data indicate mean of triplicate values for each T cell line and are representative of three independent experiments (A–B). (C–D) IFN-γ release by LCD4.G and LCD4.CD T cells in response to plate-bound soluble CD1a (sCD1a) pretreated with LppX_12mer or Ag85A_12mer peptides in the absence of LCs. Data indicate mean ± SEM of triplicate values and are representative of seven (LCD4.G) or six (LCD4.CD) independent experiments. Data were analyzed using a Friedman test followed by Dunn’s post hoc test (*P < 0.05 and **P < 0.01). See also Figure S6.

### Functional dependence on the CD1a–peptide–TCR complex

To determine whether CD1a alone is sufficient to activate T cells, we replaced LCs in the T cell assay with plate-bound soluble CD1a (**Figure 6C and 6D**). IFN-γ release by LCD4.G and LCD4.CD T cells occurred only when all three components T cells, peptide, and soluble CD1a were present. Omission of any component abolished cytokine production, an outcome that is consistent with CD1a-peptide complex recognition by T cells.

### Detection of peptide–CD1a-specific T cells in leprosy and effector cytokine production by cognate T cell lines

Fluorescent CD1a tetramers were generated by treating biotinylated sCD1a with either the LppX_12mer or Ag85A_12mer peptide, followed by cross-linking with fluorophore-conjugated streptavidin. Flow cytometric and confocal microscopy analysis demonstrated highly specific binding of each tetramer to its cognate CD1a-restricted T cell line (**Figure 7A, 7B, S7A, and S7B**). The LCD4.G T cell line bound the LppX_12mer CD1a tetramer at higher frequency (13.9 ± 1.5%), as compared to the untreated (0.42 ± 0.11%) or Ag85A_12mer (0.44 ± 0.12%) (*p* = 0.0001 for each comparison). Conversely, the LCD4.CD T cell line bound the Ag85A_12mer CD1a tetramer at high frequency (16.3 ± 0.9%) as compared to untreated (0.09 ± 0.06%) or LppX_12mer (0.10 ± 0.08%) tetramers (*p* < 0.0001 for each comparison).

**Figure 7.**
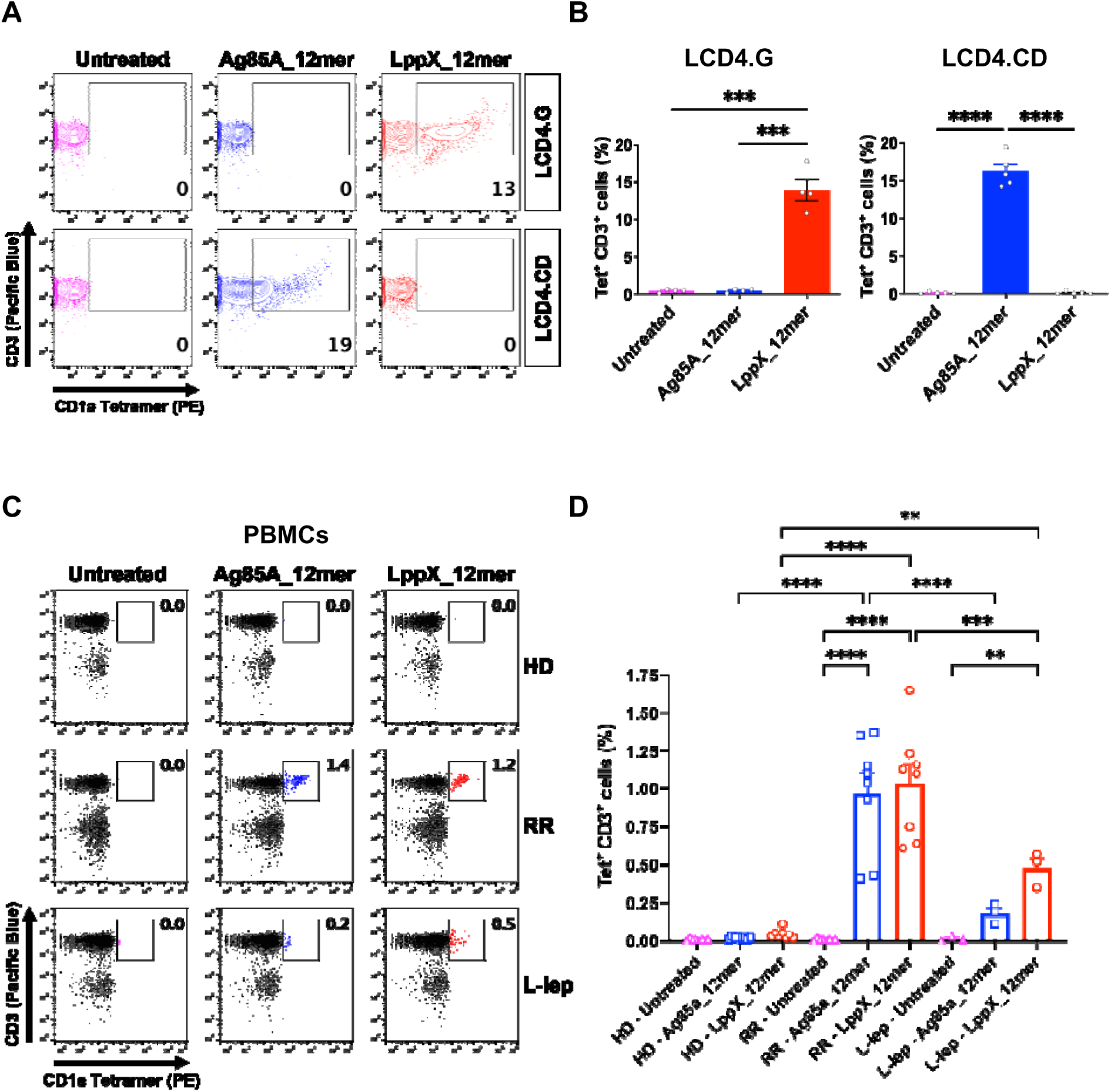
TCR recognition of CD1a–peptide complexes. (A–B) Flow cytometry analysis of LCD4.G and LCD4.CD T cell lines stained with PE-conjugated CD1a tetramers treated with Ag85A_12mer or LppX_12mer. Percent of tetramer⁺ CD3⁺ T cells are indicated. Data indicate mean ± SEM of four (LCD4.G) or five (LCD4.CD) independent experiments. Data were analyzed by one-way ANOVA followed by Tukey’s post hoc test (***P < 0.001 and ****P < 0.0001). (C–D) CD1a–peptide tetramer staining of PBMCs from healthy donors (HD), and reversal reaction (RR) and lepromatous leprosy (L-lep) patients. The percentage of tetramer⁺ CD3⁺ T cells is indicated and represents data from eight HD, eight RR, three L-lep samples. Data indicate mean ± SEM and were analyzed using a mixed-effects model (REML) estimation, followed by Tukey’s post hoc test (**P < 0.01, ***P < 0.001, and ****P < 0.0001). See also Figure S7.

We next used peptide-treated CD1a tetramers to identify CD1a–peptide–specific T cells in PBMC samples from a cohort of eleven patients with leprosy and eight healthy (**Figure 7C, 7D, and S7C**). The clinical diagnosis of leprosy was established according to the Ridley–Jopling criteria^42^. CD1a tetramers treated with either the LppX_12mer or Ag85A_12mer peptide labeled approximately 1% of CD3⁺ T cells from patients with leprosy reversal reactions (RR; n = 8) (LppX_12mer: 1.03 ± 0.12%; Ag85A_12mer: 0.97 ± 0.13%), frequencies that were significantly greater than the near-background levels observed in healthy donors (HD; n = 8) (LppX_12mer: 0.04 ± 0.01%; Ag85A_12mer: 0.02 ± 0.01%, *p* < 0.0001 for both). In patients with lepromatous leprosy (L-lep; n = 3), CD1a–peptide tetramer⁺ T cells were detected at intermediate frequencies (LppX_12mer: 0.48 ± 0.07%; Ag85A_12mer: 0.18 ± 0.04%), which were significantly higher than HD for the LppX_12mer (*p* = 0.0057) but not for the Ag85A_12mer, and significantly lower than RR patients for both peptides (LppX_12mer: *p* = 0.0009; Ag85A_12mer: *p* < 0.0001). Control tetramers lacking peptide yielded similarly low frequencies of tetramer⁺ T cells across all groups (HD: <0.01%; RR: <0.01%; L-lep: 0.01 ± 0.01%).

To determine whether the cognate CD1a-restricted T cell lines exhibited effector functions consistent with antimicrobial activity, we measured Th1- and Th17-associated cytokines following anti-CD3/CD28 stimulation (**Figure S7D and S7E**). Both LCD4.G and LCD4.CD exhibited robust cytokine secretion profiles characterized by particularly high levels of IL-26 production. LCD4.G released approximately 90 ng/ml of IL-26 and 30 ng/ml of IFN-γ, while LCD4.CD produced approximately 100 ng/ml of IL-26 and 40 ng/ml of IFN-γ. Both cytokines have been implicated in antimicrobial defense against mycobacteria, including macrophage activation and direct antibacterial activity^28,31,43^. Together, these findings demonstrate that peptide–CD1a–restricted T cells are more readily detectable in patients with relative resistance to leprosy and exhibit antimicrobial effector protein expression compatible with a role in host defense against *M. leprae*.

In summary, we identified two microbial peptide antigens (LppX_12mer and Ag85A_12mer) recognized by CD1a-restricted T cells, provided biochemical and biophysical evidence that these peptides bind to CD1a, and demonstrated that antigen-specific TCRs recognize and respond to these complexes in a CD1a–peptide–TCR–dependent manner.

Together, these findings demonstrate that CD1a can present canonical microbial peptides as part of a cell-mediated immune response in leprosy, extending the known spectrum of CD1a ligands beyond non-canonical lipopeptides and small hydrophobic molecules.

## DISCUSSION

Our findings expand the emerging paradigm of CD1a antigen presentation by demonstrating that canonical, non-lipidated microbial peptides, produced by standard protein degradation, bind CD1a and are recognized by T cells. These outcomes challenge the long-standing view that CD1-restricted T cells recognize only lipid-containing antigens or chemically diverse, non-peptidic hydrophobic ligands. By identifying canonical peptide epitopes from *M. leprae* and *M. tuberculosis* proteins (LppX and Ag85A), we show that the boundary between MHC-mediated peptide presentation and CD1-mediated lipid presentation is more flexible than previously recognized. In this respect, CD1a can resemble MHC molecules functionally by presenting protein-derived fragments to T cells, although the structural basis and binding topology remain to be determined. Early studies in mice demonstrated that CD1d molecules could present peptide antigens to T cells^44–46^. However, prior to our findings, no specific microbial peptide had been identified as a cognate ligand for any CD1 isoform. Our data provide new evidence that canonical, non-lipidated microbial peptides can be presented by CD1 molecules and recognized by T cells.

We assembled multiple layers of evidence to support this model: demonstrating binding of microbial peptides to CD1a altered its electrophoretic mobility, and SPR revealed a *K*_D_ of ∼75 μM for the Ag85A_12mer peptide. Compared to previously described hybrid antigens like dideoxymycobactin and synthetic acylated form of the HIV NEF protein^33^, which both contain a lipid tail that was shown to be required for anchoring to CD1a and CD1c, respectively, and peptidic branches for TCR contact, our canonical peptides represent a structurally distinct class that lack lipid tails^2,14^. The mycobacterial peptides rely on internal hydrophobic amino acids to engage CD1a, suggesting that CD1a may recognize peptides via mechanisms that partially mimic lipid binding, but without requiring a covalently attached, distinct lipid anchor. Further studies are required to determine where these peptides bind on the CD1a surface.

We showed that these peptide–CD1a complexes are specifically recognized by T cell receptors. CD1a–peptide tetramers stained the cognate T cells (but not unrelated T cells), providing evidence of TCR binding to the peptide–CD1a complex. Furthermore, addition of T cells to plate bound soluble CD1a and peptide markedly induced IFN-γ release in the absence of APCs, demonstrating that TCR engagement of the peptide–CD1a complex is required for T cell activation. In additional functional studies, transferring the LCD4.G TCR into a TCR-deficient 5KC T cell hybridoma conferred a CD1a-restricted response to the LppX peptide, demonstrating that the TCR alone is sufficient to endow peptide–CD1a specificity. Together, these results confirm that the peptide–CD1a complex is the bona fide antigenic determinant recognized by these T cells.

We identified the antigenic molecules as protein-derived peptides through biochemical purification, mass spectrometry, recombinant protein validation, and synthetic peptide mapping. Each T cell line’s activity was tracked to protease-sensitive fractions, and ultimately a single 12-amino-acid peptide epitope was pinpointed for each line. The discovery of these precise peptide sequences (from LppX and Ag85A) provides strong evidence that these T cells are responding to peptide epitopes. In summary, we have provided biochemical, biophysical, and functional evidence that CD1a presents canonical microbial peptides to T cells, extending the universe of CD1a antigens. This extends beyond our previous findings that CD1a-restricted T cell lines recognize *M. leprae* nonpeptide antigens^32^.

We show that CD1a binds these peptides by multiple criteria; however, the structural basis of this interaction remains unresolved. Several models are possible. First, the peptides may partially occupy the CD1a binding groove, supported by the presence of hydrophobic residues and an approximate correspondence between peptide length and the dimensions of the CD1a groove^13,47^. This possibility is consistent with the precedent of dideoxymycobactin, a non-ribosomal peptide-like scaffold covalently linked to alkyl chains that is crystallographically anchored within the CD1a groove^2,14^. Second, peptides may associate with alternative regions of CD1a, including the A′ pocket in an antigen-independent manner^48^. Third, peptides may be presented through noncanonical modes of interaction, including sideways recognition, as described for CD1a^49^ and CD1c^50^. Finally, it is possible that endogenous lipids contribute to stabilization of the complex, allowing co-presentation of peptide and lipid ligands. These possibilities are consistent with prior studies showing that CD1a can bind chemically diverse non-lipid ligands, including small hydrophobic molecules such as benzyl benzoate and farnesol^18^ and the urushiol congener C15:2^16^, and that CD1d can present nonlipid small molecules to T cells^51^. Further studies will be required to define the structural basis of peptide–CD1a interaction and its relationship to T cell receptor recognition.

Because CD1a exhibits very limited coding polymorphism relative to HLA molecules^52^, CD1a-presented peptide antigens would be expected to be less constrained by host antigen-presenting genotype than conventional HLA-presented peptides. However, regulatory variation can influence CD1a expression in cultured DCs, indicating that the prevalence and magnitude of CD1a-restricted responses may vary among individuals^53^. Thus, CD1a-presented peptides could support HLA-independent approaches to immune monitoring or vaccine design, although population-level prevalence and protective function remain to be established. In this context, the mycobacterial peptides identified here from LppX and Ag85A represent candidate antigens for stimulating or tracking CD1a-restricted T cell responses. Notably, Ag85A is already a component of several TB vaccine candidates as a dominant antigen for conventional T cells, and our findings reveal that it also contains a CD1a-restricted peptide epitope. Similarly, an LppX-based protein vaccine also induced a strong IFN-γ response in mice^54^, raising the possibility that MHC- and CD1a-restricted mechanisms could both be relevant to vaccine design. CD1a–peptide tetramers may also provide tools to detect and quantify these T cells in blood or tissues as potential biomarkers of mycobacterial immunity.

The presence of peptide–CD1a–specific T cells in leprosy patients with reversal reactions indicates that such responses arise in vivo. Supporting their in vivo relevance, CD1a–peptide tetramers identified LppX and Ag85A specific T cells at frequencies of ∼1% of circulating CD3⁺ T cells in patients undergoing reversal reactions that are actively controlling the infection, 2-5-fold greater than in lepromatous patients, those with progressive disease, and >25-fold greater than in healthy donors. The CD1a-restricted T cell lines characterized here produced IFN-γ in response to cognate peptide–CD1a stimulation and secreted both IFN-γ and IL-26 following polyclonal anti-CD3/CD28 stimulation. IFN-γ is a central mediator of macrophage activation and vitamin D–dependent antimicrobial pathways^27,28,31,55^, while IL-26 has been shown to exert direct antibacterial effects and modulate innate responses at epithelial and myeloid surfaces^43^. Previously, we showed that the CD1a-restricted T cell line LCD4.G could mediate an antimicrobial response against *M. leprae* in Langerhans cells through the production of IFN-γ, which induced autophagy and cathelicidin expression in infected DCs^31^. However, caution is warranted in assigning functional significance given the small number of T cell lines studied and that in vitro cytokine release does not equate to protection.

In conclusion, this study provides evidence that human CD1a can present canonical microbial peptides to T cells, revising the long-held view that CD1 molecules are restricted to lipids and hydrophobic small molecules. The CD1a-restricted T cells recognizing these peptides exhibit antimicrobial functions consistent with host defense, particularly at the skin barrier.

These findings broaden our understanding of T-cell antigen recognition, unveiling a previously unappreciated facet of the immune system that might now be leveraged for translational benefit.

## METHODS

### Patients

Leprosy patients were recruited at the Los Angeles County/USC Medical Center and at Fiocruz in Brazil under IRB-approved protocols (**Table S3**). Blood samples from healthy donors were obtained in accordance with protocols approved by the institutional review board at University of California, Los Angeles (#11-001274). All donors provided written informed consent for the collection of peripheral blood and subsequent analysis.

### T cell enriched media

For all in vitro T cell assays, unless otherwise indicated, cells were cultured in T cell–enriched medium (TEM) prepared by supplementing RPMI-1640 with L-glutamine (Gibco, catalog no. 11875-093) with 55 μM 2-mercaptoethanol (Gibco, catalog no. 21985-023), 10 mM HEPES (Gibco, catalog no. 15630-080), 1 mM sodium pyruvate (Gibco, catalog no. 11360-070), MEM non-essential amino acids diluted to 0.5x (Gibco, catalog no. 11140-050), MEM essential amino acids diluted to 0.25x (Gibco, catalog no. 11130-051), Penicillin-Streptomycin-Glutamine solution at 1x (Gibco, catalog no. 10378-016), and 10 μg/ml gentamicin (Gibco, catalog no. 15710-064). All components were added to the RPMI base medium and sterile-filtered through a 0.2 μm filter prior to use.

### Isolation of T lymphocytes from human skin and T cell lines

Skin biopsy specimens were trimmed to remove epidermis and subcutaneous fat, then placed in a tissue sieve fitted with a 64-µm mesh (Bellco Glass Inc.) over a petri dish containing RPMI 1640 supplemented with 10% HyClone FBS (Cytiva). The tissue was minced into ∼1 mm³ fragments using sterile scalpels and mechanically dissociated by gently pressing through the mesh with a flattened glass rod to obtain a single-cell suspension. Lymphocytes were enriched by Ficoll-Paque Plus (Cytiva) density gradient centrifugation, followed by two washes in RPMI with 10% FBS.

T cell lines were generated from leprosy RR patient skin biopsy samples by seeding T cells (100–10,000 cells/well) into 96-well round-bottom plates containing TEM supplemented with 10% allogeneic human serum and recombinant IL-2 (1 nM; Chiron or Peprotech).

Allogeneic CD1a⁺ LCDCs (10,000 cells/well) were added together with antigen (5 µg/ml sonicated *M. leprae*), and cultures were maintained for 2–3 weeks with IL-2–containing medium replenished every 2–3 days. LCD4.C, LCD4.D, and LCD4.G CD4^+^αβ T cell lines identities were confirmed by TCR sequencing.

During the second and third weeks of co-culture, T cell–LCDC co-cultures were harvested prior to each experiment. Cells were resuspended in MACS buffer supplemented with 2% FBS and subjected to magnetic labeling with anti-CD1a (CD1a MicroBeads, 130-051-001; Miltenyi Biotec), anti-CD8 (CD8 MicroBeads, 130-045-201; Miltenyi Biotec), and anti-γ/δ T cell microbeads (130-050-701; Miltenyi Biotec). Residual LCDCs, CD8αβ T cells, and γδ T cells were depleted by positive magnetic selection according to the manufacturer’s instructions. The purity of the resulting T cell lines was assessed by flow cytometry prior to each experiment.

### Generation of CD1-expressing LC-like DCs (LCDCs)

Langerhans cell-like dendritic cells (LCDCs) were derived in vitro from human Cord Blood CD34^+^ Progenitor Cells (Lonza 2C-101A) as previously described^31,32,34^. CD34^+^ hematopoietic progenitors were cultured in RPMI 1640 medium in the presence of SCF (25 ng/ml), GM-CSF (100 ng/ml), and TNF (500 ng/ml). At day 5, the remaining cells were incubated in the presence of GM-CSF (100 ng/ml) and TGF-β1 (1 ng/ml) to increase CD1a expression.

LCDCs were harvested at day 10 to 12, yielding CD1a^+^CD207^+^ (langerin^+^) dendritic cells resembling epidermal LCs. Of the CD1a^+^ LCDCs, 80-90% were found to also be CD207^+^, by surface staining. Cells were irradiated (50 Gy) before use and served as APCs.

### TCR sequencing

T cell receptor (TCR) α and β chain sequences were determined from established T cell lines. Cell pellets were harvested and immediately preserved in RNAlater solution (Ambion) until processing. Total RNA was extracted using the RNeasy plus mini kit (Qiagen). RNA concentrations were then quantified using a Qubit Fluorometer and RNA integrity was assessed using the Agilent TapeStation (Agilent). Samples with RIN (RNA integrity number) ≥ 8 were used for this study.

Total RNA was reverse-transcribed into cDNA using template switch oligonucleotide (TSO), primers for human TCR constant regions and SMARTScribe reverse transcriptases (Clontech) according to the manufacturer’s instructions. TCRα and TCRβ transcripts were amplified by two subsequent PCR using nested primers for the constant gene region. All libraries from purified TCR amplicons were prepared using the Nextera XT Index kit (Illumina Inc.) according to the manufacturer’s instructions for the “16S Metagenomic Sequencing Library Preparation” protocol with minor modifications. Sequencing was performed on an Illumina MiSeq® sequencer using the 600-cycle MiSeq Reagent Kit v3 (Illumina) with paired-end reads. Raw sequencing reads were processed for FASTQ conversion and demultiplexing using MiSeq Reporter. MiXCR software were used to extract TCR CDR3 sequences from sequencing data. All gene names used are according to IMGT nomenclature. TCR repertoire comparison was done in R (version 3.1.2, http://www.R-project.org/). The IMGT/Junction Analysis tool was used to analyze in detail the iCDR3 amino acid sequences and CDR3 V-D-J junctions. Paired TCRα and TCRβ sequences were determined for each T cell line and used for downstream analyses. TCR sequencing data were deposited in the NCBI BioProject database under accession number PRJNA1455222.

### Isolation of human peripheral blood mononuclear cells (PBMCs) and handling

Peripheral blood mononuclear cells (PBMCs) were obtained from the peripheral blood of healthy individuals employing a Ficoll-Paque Plus (Cytiva) density sedimentation gradient in an endotoxin-free environment. The isolated PBMCs underwent lysis of red blood cells using an ACK lysis buffer (Thermo Fisher). PBMCs were then resuspended in cryopreservation medium containing 10% DMSO and stored in liquid nitrogen until further analysis.

### Antigen Preparation and Presentation Assays

Bacterial sonicates or lysates were prepared by disruption of cells via probe sonication or French press in phosphate-buffered saline (PBS). *M*. *tuberculosis* sonicate, *M. smegmatis* lysate, mycolic acid, manLAM, MLSA, MLMA, MLCwA, and CWC, as well as *E. coli* sonicate, and *P. acnes* sonicate, were provided by Dr. John Belisle (Colorado State University). Assays with *S. aureus* Ag^−^ sonicate were performed using the triple-mutant strain DU5938 (Hla^−^ Hlb^−^ Hlg^−^)^56^. *M. leprae* sonicate and total lipid extracts were obtained from BEI Resources (NIAID/NIH). C80-86 GMM^4^ was purified from *M. phlei* (Tan-Yun Cheng, Brigham and Women’s Hospital). Dideoxymycobactin^3^ was synthesized by David C. Young (Brigham and Women’s Hospital), and C35 mannosyl phosphodolichol^7^ was synthesized by Gurdyal S. Besra (University of Birmingham).

For antigen presentation assays, purified T cells (1×10^5^ per well) were co-cultured with APCs (either 5×10^4^ or 1×10^5^ CD1a^+^ LCDCs) in 200 μL TEM with 10% human serum in 96-well plates. Soluble antigens (bacterial lysates, fractions, or proteins) were pre-pulsed onto APCs at the indicated concentrations (typically ∼10 μg/ml for lysates, or equimolar protein amounts) and incubated for 1 h at 37°C before T cells were added.

### T cell activation and cytokine quantification

In some experiments, T cell lines were stimulated with anti-CD3 and anti-CD28 monospecific antibody complexes (ImmunoCult, STEMCELL Technologies) to induce polyclonal activation. Cells were incubated at 37°C in TEM supplemented with 10% human serum, and culture supernatants were collected after 48 h of stimulation. Cytokine secretion was quantified by enzyme-linked immunosorbent assay (ELISA) for IL-26 (Cusabio Biotech) and for IL-17, IL-22, IL-4, IL-10 (DuoSet kits; R&D Systems), and IFN-γ (BD Biosciences), according to the manufacturers’ instructions. Cytokine concentrations were determined by comparison with recombinant cytokine standard curves and are reported as ng/ml.

### IFN-**γ** release and lymphocyte proliferation assays

T cell proliferation was assessed by [³H]-thymidine incorporation (ICN Biomedicals Inc.; 1 μCi per well added during the final 4 h of a 3-day culture) and expressed as counts per minute (CPM). Cells were harvested onto 96-well microfiber filters (Filtermat A, Revvity) using a Tomtec Mach III harvester, and liquid scintillation counts were measured using a PerkinElmer/Wallac 1450 MicroBeta TriLux. IFN-γ production in matching cultures (derived from the same experimental wells) was measured by ELISA from 24 h culture supernatants and reported as pg/ml. All assays were performed in triplicate, and data are presented as mean ± SEM of replicate wells.

### Functional inhibition of CD1a using neutralizing antibodies

CD1a-mediated blocking of *M. tuberculosis* -specific immune responses was assessed using IFN-γ release and lymphocyte proliferation assays in parallel. CD1a^+^ LCDCs were cultured with the indicated T cell lines in TEM medium alone or in the presence of *M. tuberculosis* sonicate, *M. tuberculosis* plus αCD1a antibody, or *M. tuberculosis* plus IgG1 isotype control antibody. Purified anti-CD1a monoclonal antibody^4^ was used at 10 µg/ml to block recognition of antigen presented on the CD1a α1–α2 platform. Isotype-matched IgG1 (clone P3.6.2.8.1) served as a control. Antibodies were added to antigen-presenting cells (APCs) 30 min prior to antigen addition and were maintained throughout the culture period.

For both assays, raw values were background-subtracted by subtracting the medium-only condition from each experimental condition. Background-corrected responses in the presence of antibody were normalized to the *M. tuberculosis*–only control condition, which was set to 100%^57^. Percent inhibition (blocking) was then calculated for the CD1a condition, with the IgG1 condition serving as a negative control. This approach enabled normalization of antigen-specific responses and comparison of antibody effects across experiments using different LCDC donors and T cell lines. The equation below summarizes the calculation of percent functional inhibition (where Ab represents either CD1a or IgG1) and is reported as the percentage of the *M. tuberculosis* relative response remaining.

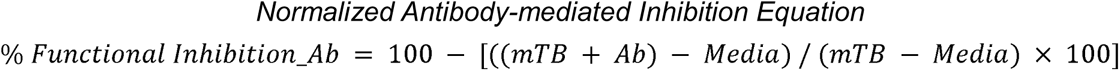

### Bacterial fractionation

*M. tuberculosis* strain H37Rv was cultured in glycerol alanine salts medium^58^. Whole cell lysate and subcellular fractions of *M. tuberculosis* were generated as previously described^59^. *M. leprae* (strains NHDP-63 and NHDP-98) and subcellular fractions of this bacterium were obtained through BEI Resources (NIAID/NIH). Cell envelope proteins of *M. tuberculosis* were extracted and isolated from the cell wall fraction by Triton X-114 phase separation as described^36,60^. The detergent phase was collected and Triton X-114 was removed by cold acetone precipitation. The protein pellet was solubilized in 10 mM ammonium bicarbonate and the protein concentration determined by the Bicinchoninic acid assay.

### Protease treatment

Aliquots of the *M. tuberculosis* Triton X-114 extract were prepared under four conditions before addition to the T cell assay: (1) untreated extract; (2) extract incubated with proteinase K (50 μg/ml for 1 h at 37°C) followed by heat inactivation at 95°C for 10 min; (3) extract heated at 95°C for 10 min in the absence of proteinase K; and (4) extract incubated with proteinase K that had been pre-inactivated by heating at 95°C for 10 min. The resulting fractions were then added to the T cell proliferation assay.

### T cell antigen identification

The *M. tuberculosis* (H37Rv) Triton X-114 extracted proteins (50 mg) were lyophilized with 7.25 M PlusOne urea (Fisher), 0.4% 3-10 ZOOM carrier ampholytes (ThermoFisher), 1.6% 4-7 pharmalytes, 1% N-octylthioglucoside (EMD Chemicals, Inc) and 2 mM DL-Dithiothreitol (DTT, Fisher). The proteins were fractioned by liquid phase isoelectric focusing using the Rotofor (Biorad) Isoelectric Focusing Preparative Cell. This yielded 20 fractions that were dialyzed against 10 mM ammonium bicarbonate using 3.5 kDa MWCO membrane. Each fraction was analyzed by SDS-PAGE and silver staining to visualize protein distribution, and tested for T cell reactivity. Active fractions were pooled and further screened with the T cell “Western blot” assay^37^. Specifically, proteins were resolved on a NuPAGE 4-12% Bis-Tris gels (ThermoFisher) and electroblotted to nitrocellulose membranes. Protein lanes on the nitrocellulose blots were horizontally cut into 30 slices of 2.5 mm. Strips of membrane containing the separated proteins from each fraction were incubated overnight with CD1a^+^ LCDCs (as APCs) and T cells (either LCD4.G or LCD4.CD) in culture medium, under conditions analogous to an ELISPOT assay. After ∼36–48 h of incubation, the membrane strips were removed, and the cells were analyzed for IFN-γ production. Alternatively, supernatants from each membrane–cell co-culture were collected and assayed by ELISA for IFN-γ, to identify which membrane strip (hence which fraction and molecular weight range) had triggered T cell activation.

To identify potential T cell reactive proteins, identical nitrocellulose strips were digested with trypsin overnight at 37°C as previously described^61^. Trypsin digested samples were dried in a vacuum centrifuge, and the nitrocellulose was dissolved with acetone at room temperature for 30 min. Precipitated peptides were collected by centrifuging at 14,000 x g for 10 minutes and then washed twice with acetone and dried at room temperature. Peptides were cleaned using PepClean C18 spin columns (Pierce) prior to LC-MS/MS.

### Mass spectrometry analyses

Peptides were concentrated using an on-line enrichment column (Thermo Scientific 5μm, 100 μm ID x 2cm C18 column). Subsequent chromatographic separation was performed on a reverse phase nanospray column (Thermo Scientific EASYnano-LC, 3μm, 75 μm ID x 100mm C18 column) using a 90 min linear gradient from 10%-30% buffer B (100% ACN, 0.1% formic acid) at a flow rate of 400 nL/min. Peptides were eluted directly into a Thermo Scientific Orbitrap Velos mass spectrometer and spectra collected over a m/z range of 400-2000 Da using a dynamic exclusion limit of 2 MS/MS spectra of a given peptide mass for 30 s (exclusion duration of 90 s). The instrument was operated in Orbitrap-LTQ mode where precursor measurements were acquired in the orbitrap (60,000 resolution) and MS/MS spectra (top 20) were acquired in the LTQ ion trap with a normalized collision energy of 35%. Compound lists of the resulting spectra were generated using Xcalibur 2.2 software (Thermo Scientific) with a S/N threshold of 1.5 and 1 scan/group.

Tandem mass spectra were extracted, charge states deconvoluted and deisotoped by ProteoWizard (MS-Convert version 3.0.). MS/MS data were analyzed using Mascot (Matrix Science, London, UK; version 2.3.02 with the Tuberculist_R27_March 2013_rev100214 database (v27 (March 2013), 8062 entries). Mascot was searched with a fragment ion mass tolerance of 0.80 Da and a parent ion tolerance of 10.0 PPM. Oxidation of methionine and carbamidomethyl of cysteine were specified in Mascot as variable modifications.

Scaffold (version Scaffold_4.3.2, Proteome Software Inc., Portland, OR) was used to validate MS/MS based peptide and protein identifications. Peptide identifications were accepted if they had > 6.0% probability to achieve an FDR < 0.5%. Protein identifications were accepted if they had > 97.0% probability to achieve < 1.0% FDR and contained at least 2 identified peptides. Protein probabilities were assigned by the Protein Prophet algorithm^62^.

### Recombinant antigen production

Recombinant C-terminal 6×His tag constructs of *M. tuberculosis* LprG, SodC, LprF, LpqZ, and BlaC were created by PCR amplification of the genes, cloning into the mycobacterial shuttle vector pVV16 and expression in *M. smegmatis*. Recombinant C-terminal 6×His tag constructs of *M. tuberculosis* LppX, Ag85A, Ag85B and Ag85C were also produced in *E. coli* BL21(DE3) using the pET expression vector. His-tagged proteins were purified from cell lysates by Ni-NTA affinity chromatography. Purified proteins were tested for endotoxin (<0.1 EU/µg). Protein identities were confirmed by mass spectrometry (peptide mass fingerprinting and intact mass measurement).

### Generation of glutathione S-transferase (GST) fusion fragments

For epitope mapping, overlapping fragments of the *lppX* or *ag85A* genes were generated by PCR, ligated with pGEMTeasy vector, and transformed into *E. coli* Top10 strain for plasmid amplification. The *lppX* and *ag85a* genes were digested from the cloning vector using defined restrictions sites and cloned in-frame into the pGEX-6P-1 vector to express GST fusion proteins covering different regions of LppX and Ag85A. These fusion proteins were affinity-purified on glutathione-Sepharose, analyzed by Western blot, and used in T cell assays at equimolar concentrations (normalized for fusion protein size).

### Synthetic peptides

Peptides were synthesized by standard Fmoc solid-phase synthesis (GenScript or MIMOTOPES). Overlapping peptides (15- to 20-mers with 5-aa overlaps) spanning regions of interest in LppX and Ag85A were initially tested (at ∼5 μg/ml) to identify reactive segments.

Once a reactive peptide was identified, truncated derivatives were synthesized to find the minimal epitope. The LppX_12mer (SHHLVRASIDLG) and Ag85A_12mer (AGCQTYKWETFL) were each synthesized at >98% purity, with a free N-terminus (no acetylation) and C-terminal free acid, to mimic a natural peptide’s ends. Biotinylated versions of these peptides (with biotin attached via an N-terminal linker) were also prepared for SPR binding studies. Peptides were dissolved in PBS and stored at -20 °C. For T cell assays, peptides were added to APC–T cell co-cultures by pre-pulsing the APCs with peptide for 1 h at 37°C. Dose–response assays were performed over a range from 0.1 μg/ml up to 10 μg/ml of peptide.

### Hydropathy Analysis

Hydropathy profiles were generated for the Ag85A_12mer (AGCQTYKWETFL) and LppX_12mer (SHHLVRASIDLG) peptides using the Kyte–Doolittle scale. Mean hydropathy values were calculated across the peptide length using a sliding window approach. Hydropathy scores were visualized as bar plots generated in Python (v3.x) using the Matplotlib library on a white background with standardized color coding. All analyses and figures were produced programmatically to ensure reproducibility.

### Mycobacterial homolog identification and comparative protein sequence analysis

To identify Ag85 homologs in *M. smegmatis*, candidate genes annotated as Ag85 family members were first retrieved from MycoBrowser (https://mycobrowser.epfl.ch/). These sequences were subsequently subjected to Protein BLAST analysis (https://blast.ncbi.nlm.nih.gov/) against *M. tuberculosis* H37Rv using the Model Organism (Landmark) database.

Homolog selection was based on percentage amino acid identity, percentage amino acid similarity, and the number of identical residues corresponding to the Ag85A peptide region of interest. Based on these criteria, two *M. smegmatis* candidates, MycoBrowser #6398 (Ag85A) and #2078 (Ag85C; annotated as Ag85B in UniProt), were initially identified for comparison with *M. tuberculosis* Ag85A (#004032) and *M. leprae* Ag85A/fbpA (#ML0097). For subsequent alignment analyses, #2078 was selected because its sequence is annotated in UniProtKB/Swiss-Prot (reviewed; ‘sp’), a manually curated, high-confidence database, whereas #6398 is annotated in UniProtKB/TrEMBL (unreviewed; ‘tr’), which contains computationally predicted entries pending curation. Accordingly, a reviewed Swiss-Prot entry was prioritized to ensure greater annotation reliability and sequence accuracy.

Protein sequence alignments of Ag85 and LppX across mycobacterial species were performed using the UniProt alignment tool^63^. Amino acid conservation and biochemical differences were assessed using the ‘physical properties’ option with hydrophobicity selected as the comparison parameter.

### Generation and Functional Assessment of Human TCR-Expressing 5KC Reporter Cells

To assess antigen-specific TCR function in a controlled cellular system, 5KC cells, murine T-cell hybridomas lacking endogenous TCR α- and β-chains were engineered to stably express human CD4, which enhances TCR–antigen interactions, as previously described^64^.

These cells were subsequently transfected with the LCD4.G T-cell receptor using a hybrid human/murine TCR α/β construct. Co-expression of CD4 and TCR was confirmed by flow cytometry. Dead cells were excluded using Live/Dead Blue dye (Invitrogen #L23105). Cells were stained with the following antibodies: FITC anti-human CD4 (clone RPA-T4, BioLegend #300506) or FITC mouse IgG1κ isotype control (clone MOPC-21, BioLegend #400110); APC anti-mouse CD3ε (clone 145-2C11, BioLegend #100312) or APC Armenian hamster IgG isotype control (clone HTK888, BioLegend #400912); and PE anti-mouse TCRβ chain (clone H57-597,

BioLegend #109208) or PE Armenian hamster IgG isotype control (clone HTK888, BioLegend #400908). TCR activation was evaluated by co-culturing 5KC-hCD4-TCR transfectants with CD1a^+^ LCDCs in the presence of *M. leprae* sonicate, recombinant LppX(-) produced in *E. coli*, or the LppX_12mer peptide. Mouse IL-2 production (DuoSet, R&D Systems), measured by ELISA, was used as the readout for TCR activation. As a positive control, LCD4.G T cells were stimulated with the same antigens, and human IFN-γ production was quantified by ELISA.

### Soluble CD1a Production

Biotinylated CD1a monomers (97–99% biotinylation), expressed in human HEK293T cells, were obtained from the NIH Tetramer Core Facility at Emory University, a major provider of tetramer reagents to the research community worldwide.

For surface plasmon resonance and isoelectric focusing gel analysis, CD1a-β2m was expressed by co-transfection of HEK293S GnTI^−^ cells as previously described^18^. Briefly, the above CD1a heavy chain and β2m gene inserts were cloned into a pHLsec vector as a single open reading frame, which included C-terminal fusion tags consisting of leucine Fos-Jun zippers, BirA biotinylation site, and a 6×His tag, with CD1a-β2m linked by a 2A linker peptide. The secreted CD1a-β2m heterodimer was purified by immobilized nickel affinity (HisTrap, Cytiva), followed by size-exclusion chromatography (Superdex 200). Expression tags and post-translational N-linked glycosylation were cleaved using Thrombin (Sigma, T6634) and Endoglycosidase H (NEB, #P0703) respectively, following manufacturer protocols, before a final purification step by size-exclusion chromatography.

### Soluble CD1a Presentation Assay

To assess the functional dependence of the CD1a–TCR complex on T cell activation, biotinylated soluble CD1a was immobilized on 96-well streptavidin-coated plates (Thermo Fisher Scientific), as previously described^17,18^, and pre-incubated with LppX_12mer or Ag85A_12mer peptides at 37°C for 1 h. Following incubation, LCD4.G and LCD4.CD T cell cultures were added to the wells. In parallel, a second plate containing T cells co-cultured with CD1a^+^ LCDCs served as a positive control. IFN-γ production in all cultures was quantified as described above.

### Isoelectric Focusing (IEF) Binding Assay

Soluble CD1a (10 μg) was incubated with LppX_12mer or Ag85A_12mer peptide (at a 20-fold molar excess) with 0.5% CHAPS-hydrate in 10 μL PBS for 16 h at room temperature to allow complex formation (for “no peptide” controls, buffer or detergent alone was added in place of the antigenic peptide). 1 ug of each sample was then subjected to analytical IEF on a precast pH 5–8 gel (Cytiva) and electrophoresis was conducted using the PhastGel system. Gels were fixed and stained with Coomassie blue for protein shift visualization. The pI position of the CD1a heavy chain band was compared between samples with and without peptide.

### Surface Plasmon Resonance (SPR)

Binding of CD1a to immobilized peptide was measured by surface plasmon resonance on a Biacore T200 instrument (Cytiva). Biotinylated LppX_12mer and Ag85A_12mer peptides were immobilized on a Series S Sensor Chip coated with Streptavidin (SA). A biotinylated DQ2.5-glia-α1a gluten peptide (PFPQPELPY)^65^ was used as a negative control. Approximately 500 response units (RU) of peptide were immobilized on the active flow cell; a reference flow cell was prepared with streptavidin alone, followed by blocking with free D-biotin. Soluble CD1a (not biotinylated, in running buffer 10 mM Tris-HCl pH 8.0, 150 mM NaCl, 2 mM EDTA) was injected over the flow cells at concentrations ranging from 0 to 150 μM (in randomized order) at 30 μL/min with a 120 s association phase, followed by dissociation for 300 s. The surface was regenerated with pulses of 2 M NaCl. All measurements were performed at 20°C and were in three independent experiments with two technical replicates each. Sensorgrams were double referenced (subtracting the reference cell signal and blank injections) and fit to a 1:1 Langmuir binding model to derive kinetic constants. Results were analyzed in BIAevaluation 4.1, and visualized using GraphPad Prism 10 (Dotmatics).

### Preparation of Peptide-treated CD1a Tetramers

CD1a tetramerization was carried out as previously described^18,48,49^. Briefly, 10 nmol of each peptide were dissolved in 45 µL PBS containing 0.5% CHAPS (Avanti Research, 850500) and incubated at 56°C for 30 min. Subsequently, 10 µg (approximately 0.2 nmol) of biotinylated CD1a monomers were added to achieve a peptide-to-monomer ratio of 50:1, followed by overnight incubation at 37°C. The next day, peptide-treated CD1 monomers were transferred to sterile tubes and tetramerized at room temperature in the dark by sequentially adding aliquots of streptavidin–phycoerythrin (PE; Thermo Fisher #S866) at 10-minute intervals. Fully assembled tetramers were stored at 4°C until use.

### CD1a Tetramer Staining of T cell lines

CD1a tetramers were validated by staining the T cell lines LCD4.G and LCD4.CD, as previously described^18,48,49^. Briefly, 1×10L T cells were incubated with human AB serum (Millipore Sigma #H4522) for 10 min, washed, and resuspended in FACS buffer (PBS containing 2 % Seradigm FBS). Cells were then stained with PE-labeled CD1a tetramer (1:25 dilution in FACS buffer) for 20 min at room temperature in the dark. Subsequently, 0.1 µg of functional-grade anti-CD3 antibody (clone OKT3, Miltenyi Biotec #130-093-387) was added per sample, and cells were incubated for 10 min at 37°C. Afterward, cells were stained with Zombie NIR viability dye (BioLegend #423105) for 20 min to exclude dead cells. Following washing, T cells were stained with Pacific Blue anti-human CD3 monoclonal antibody (clone UCHT1, BioLegend #300431) or the corresponding isotype control (clone MOPC-21, BioLegend #100151) for 25 min and fixed in 2% formaldehyde prior to analysis. Data were acquired based on FSC–SSC parameters and doublet exclusion on a SORP LSRFortessa X-20 (BD Biosciences) using FACSDiva software version 8.0.2 (BD Biosciences) at the UCLA Flow Cytometry Core Facility. Data were analyzed using FlowJo software version 10.8.2 (BD Biosciences).

In selected experiments, T cell lines were transferred onto glass slides and imaged by confocal microscopy as previously described^66^, using a Leica TCS SP8 Digital LightSheet Laser Scanning Confocal Microscope at the Advanced Light Microscopy and Spectroscopy Laboratory, California NanoSystems Institute, UCLA. NucRed 647 dye (Thermo Fisher #R37106) was used for nuclear staining prior fixation. Images were acquired using a CS2 100x/1.4 NA oil immersion objective and controlled with Leica Microsystems Application Suite X (LAS X) software.

### CD1a Tetramer Staining of PBMCs

Cryopreserved PBMCs were rapidly thawed in a 37°C water bath and incubated overnight at 37°C in complete TEM supplemented with IL-15 (1 ng/ml). PBMCs (1×10L to 5×10L) were then stained as described above for T cell lines, with minor modifications. Zombie Green viability dye (BioLegend #423111) was included to exclude nonviable cells. To exclude monocytes and B cells, anti-human CD14–PerCP/Cyanine5.5 (clone 61D3, eBioscience #45-0149-42) and anti-human CD19–PerCP/Cyanine5.5 (clone SJ25C1, BD Biosciences #340950) antibodies were added, respectively. Isotype-matched controls included mouse IgG1 κ antibodies, clones P3.6.2.8.1 (eBioscience #45-4714-82) and X40 (BD Biosciences #347212). Samples were acquired on the Attune NxT Acoustic Focusing Cytometer (Thermo Fisher Scientific) and analyzed using FlowJo software.

### Statistical Analysis

Statistical analysis and graphing were undertaken with GraphPad Prism software version 10. Statistics reported are of entire series of experiments and described as mean ± the standard error (SEM). For comparison between three or more groups with normally distributed, matched or repeated data, we used repeated measures one-way ANOVA with Tukey’s multiple comparisons test with individual variances computed for each comparison. For comparison between three or more groups with non-normal, matched or repeated data, we used Friedman test with Dunn’s multiple comparisons test. For datasets containing missing values (e.g., unequal numbers of donors across groups), a mixed-effects model using restricted maximum likelihood (REML) estimation was applied, followed by Tukey’s multiple comparisons test. A P value < 0.05 was considered statistically significant.

### Limitations of the study

This study has several limitations. First, although biochemical and functional data support direct association of microbial peptides with CD1a, the structural basis of peptide presentation remains unresolved. In particular, we have not determined whether these peptides occupy the canonical CD1a lipid-binding cleft, bind elsewhere on the CD1a surface, or are co-presented with endogenous lipids that stabilize the complex. Second, identification of the Ag85A epitope relied on a limited set of hypothesis-guided candidate peptides rather than exhaustive tiling of the entire N-terminal fragment, and additional CD1a-presented sequences cannot be excluded. Third, TCR transfer experiments were performed for the LppX-specific LCD4.G TCR but not for the Ag85A-specific LCD4.CD TCR. Fourth, the patient cohort used for tetramer analysis was limited in size, particularly for lepromatous leprosy, and tetramer-positive PBMCs were not fully phenotyped or directly tested for antigen-induced effector function. In addition, tetramer-positive cells in PBMCs were quantified within the CD3^+^ compartment, as is common in tetramer-based analyses; however, their precise lineage (e.g., CD4^+^, CD8^+^, γδ), differentiation state, and antigen-induced effector functions were not further resolved in this study. Fifth, although the identified T cell lines produce IFN-γ and IL-26, in vitro cytokine production does not establish protective function in vivo. Finally, although CD1a is nearly monomorphic, the population breadth and protective capacity of CD1a–peptide-specific T cell responses remain to be established in larger and clinically diverse cohorts.

## RESOURCE AVAILABILITY

### Lead contact

Further information and requests for reagents should be directed to the lead contact, Robert L. Modlin.

### Materials availability

No additional unique reagents, beyond the cell lines and TCR constructs described above, were generated for this study.

### Data and code availability

TCR sequencing data generated in this study have been deposited in the NCBI BioProject database under accession number PRJNA1455222 and will be released upon publication. This paper does not report new original codes. Any additional information required to reanalyze the data reported in this paper is available from the lead contact upon request.

## ACKNOWLEDGMENTS

We thank the NIH Tetramer Core Facility (NIH contract 75N93020D00005 and RRID:SCR_026557) for providing CD1a monomers; the UCLA Flow Cytometry Core Facility for assistance with flow cytometry; the California NanoSystems Institute, Advanced Light Microscopy Core Facility for assistance with the confocal studies; Barry R. Bloom for helpful discussions; and Tan-Yun Cheng and David C. Young (Brigham and Women’s Hospital) for providing CD1 antigens.

This work was supported by NIH grants R01 AI022553, R01 AR040312, R01 AI166313, and P50 AR080594 for RM; NHMRC Emerging Leadership Investigator Grant (2027104) for AS; ARC Discovery grant (DP260100848) and NHMRC investigator award (APP2008981) to JR. R01AR048632, R01 AI049313, and Wellcome Trust Discovery Award to DBM.

## AUTHOR CONTRIBUTIONS

**Conceptualization:** B.J.d.A.S., A.d.J., D.M.Z., D.B.M., K.R.N., E.M., A.Se., J.R., J.T.B., R.L.M.; **Methodology:** B.J.d.A.S., A.d.J., L.A.F., M.A.M.M., A.L., A.Sh., J.K., P.A.S., A.C., H.J.K., C.A.M.e.S., K.W., J.B., A.M., K.A.T., D.B.M., K.R.N., E.M., A.Se., J.R., J.T.B., R.L.M.; **Investigation:** B.J.d.A.S., A.d.J., L.A.F., M.A.M.M., A.L., A.Sh., J.K., P.A.S., A.C., H.J.K., C.A.M.e.S., K.W., J.B., P.J.B., A.M., K.A.T., E.N.S., R.O.P., K.R.N., M.T.O.; **Formal Analysis:** B.J.d.A.S., A.d.J., A.Sh., J.K., P.A.S., A.C., P.J.B., A.M., K.A.T., E.N.S., R.O.P., D.M.Z., D.B.M., K.R.N., E.M., A.Se., J.R., M.T.O., J.T.B., R.L.M.; **Data Curation:** B.J.d.A.S., A.C., A.M., K.A.T.; **Resources:** P.J.B., D.M.Z., D.B.M., E.M., A.Se., J.R., J.T.B., R.L.M.; **Clinical Samples and Classification:** E.N.S., R.O.P., M.T.O.; **Writing – Original Draft:** B.J.d.A.S., A.d.J., A.C., D.B.M., K.R.N., E.M., J.R., J.T.B., R.L.M.; **Writing – Review & Editing:** B.J.d.A.S., A.d.J., A.Sh., A.M., K.A.T., D.B.M., K.R.N., E.M., J.R., J.T.B., R.L.M.; **Supervision:** A.d.J., P.A.S., J.T.B., R.L.M.; **Funding Acquisition:** J.T.B., R.L.M.

## Declaration of generative AI and AI-assisted technologies in the writing process

During the preparation of this work the author(s) used ChatGPT (OpenAI) in order to iteratively edit and refine manuscript text through a back-and-forth process in which author-written drafts were revised and evaluated. After using this tool, the authors reviewed and edited the content as needed and take full responsibility for the content of the published article.

## SUPPLEMENTAL INFORMATION

Document S1. Figures S1–S7, and Tables S1–S3

