## Supplemental Data 1 for "CD1a-Mediated Presentation of Canonical Microbial Peptides to T Cells"

Figure S1

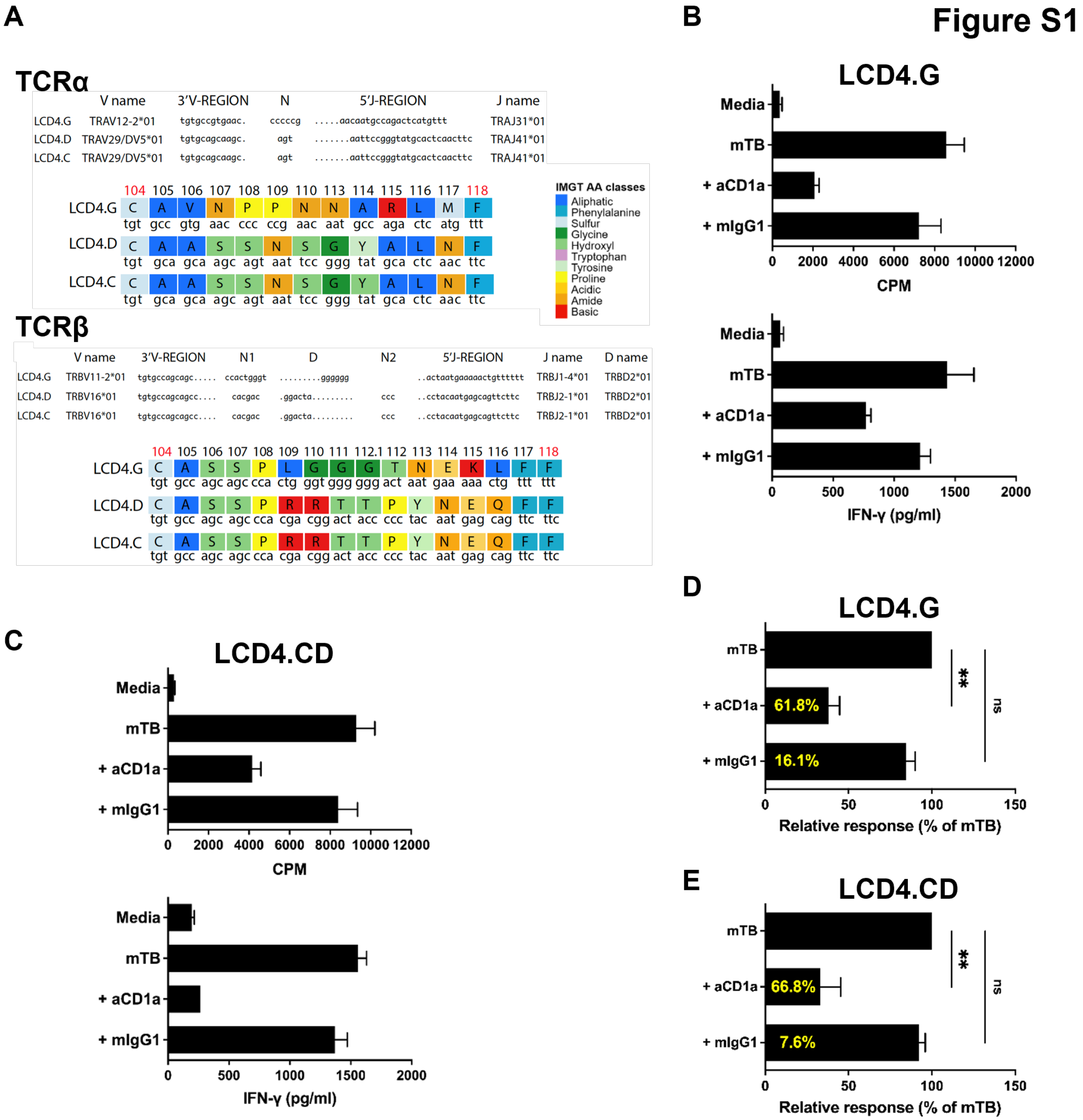

A
TCRα
V name 3'V-REGION N 5'J-REGION J name
LCD4.G TRAV12-2*01 tgtgccgtgaac. cccccg .....aacaatgccagactcatgttt TRAJ31*01
LCD4.D TRAV29/DV5*01 tgtgcagcaagc. agt .......aattccgggtatgcactcaacttc TRAJ41*01
LCD4.C TRAV29/DV5*01 tgtgcagcaagc. agt .......aattccgggtatgcactcaacttc TRAJ41*01
104 105 106 107 108 109 110 113 114 115 116 117 118
LCD4.G C A V N P P N N A R L M F
tgt gcc gtg aac ccc ccg aac aat gcc aga ctc atg ttt
LCD4.D C A A S S N S G Y A L N F
tgt gca gca agc agt aat tcc ggg tat gca ctc aac ttc
LCD4.C C A A S S N S G Y A L N F
tgt gca gca agc agt aat tcc ggg tat gca ctc aac ttc
IMGT AA classes
Aliphatic
Phenylalanine
Sulfur
Glycine
Hydroxyl
Tryptophan
Tyrosine
Proline
Acidic
Amide
Basic
TCRβ
V name 3'V-REGION N1 D N2 5'J-REGION J name D name
LCD4.G TRBV11-2*01 tgtgccagcagc..... ccactgggt .........gggggg ..actaatgaaaaactgtttttt TRBJ1-4*01 TRBD2*01
LCD4.D TRBV16*01 tgtgccagcagcc.... cacgac .ggacta......... ccc ..cctacaatgagcagttcttc TRBJ2-1*01 TRBD2*01
LCD4.C TRBV16*01 tgtgccagcagcc.... cacgac .ggacta......... ccc ..cctacaatgagcagttcttc TRBJ2-1*01 TRBD2*01
104 105 106 107 108 109 110 111 112.1 112 113 114 115 116 117 118
LCD4.G C A S S P L G G G T N E K L F F
tgt gcc agc agc cca ctg ggt ggg ggg act aat gaa aaa ctg ttt ttt
LCD4.D C A S S P R R T T P Y N E Q F F
tgt gcc agc agc cca cga cgg act acc ccc tac aat gag cag ttc ttc
LCD4.C C A S S P R R T T P Y N E Q F F
tgt gcc agc agc cca cga cgg act acc ccc tac aat gag cag ttc ttc
B
LCD4.G
Media
mTB
+ aCD1a
+ mIgG1
0 2000 4000 6000 8000 10000 12000
CPM
0 500 1000 1500 2000
IFN-γ (pg/ml)
C
LCD4.CD
Media
mTB
+ aCD1a
+ mIgG1
0 2000 4000 6000 8000 10000 12000
CPM
0 500 1000 1500 2000
IFN-γ (pg/ml)
D
LCD4.G
mTB
+ aCD1a 61.8%
+ mIgG1 16.1%
**
ns
0 50 100 150
Relative response (% of mTB)
E
LCD4.CD
mTB
+ aCD1a 66.8%
+ mIgG1 7.6%
**
ns
0 50 100 150
Relative response (% of mTB)

**Figure S1. TCR-defined clonotypic diversity of human skin T cell lines and blockade of antigen presentation by anti-CD1a monoclonal antibody.**

(A) Analysis of TCRα and TCRβ CDR3 sequences V(D)J junctions for $CD4^+$ T cell lines LCD4.G, LCD4.C, and LCD4.D. Dots in the 3'V-REGION, D-REGION and 5'J-REGION indicate nucleotides trimmed in the rearranged sequence, by comparison to the corresponding germline 3'V-REGION, D-REGION, and 5'J-REGION. N, N1, N2, indicates N-REGIONs. Each JUNCTION nucleotide sequence was translated in amino acid sequences and JUNCTION amino acids are colored according to the IMGT amino acid classes for chemical characteristics.

(B) LCD4.G T cell line proliferation and IFN-γ release in response to sonicated *M. tuberculosis* (mTB) in the presence of anti-CD1a or IgG1 isotype control.

(C) Proliferation and IFN-γ responses of the LCD4.CD T cell line to sonicated *M. tuberculosis* in the presence of anti-CD1a or IgG1 isotype control. Data indicate mean ± SEM of triplicate values and are representative of three independent experiments (B–C).

(D–E) *M. tuberculosis*–induced responses in the presence of anti-CD1a blocking antibodies or isotype control. Values shown inside the bars (yellow) indicate the percentage inhibition. Data indicate mean ± SEM and are representative of six independent experiments for LCD4.G (D) and LCD4.CD (E). Data were analyzed using a Friedman test followed by Dunn's post hoc test (**P < 0.01; ns, not statistically significant). Statistical analyses were performed on background-corrected values.

### Figure S2

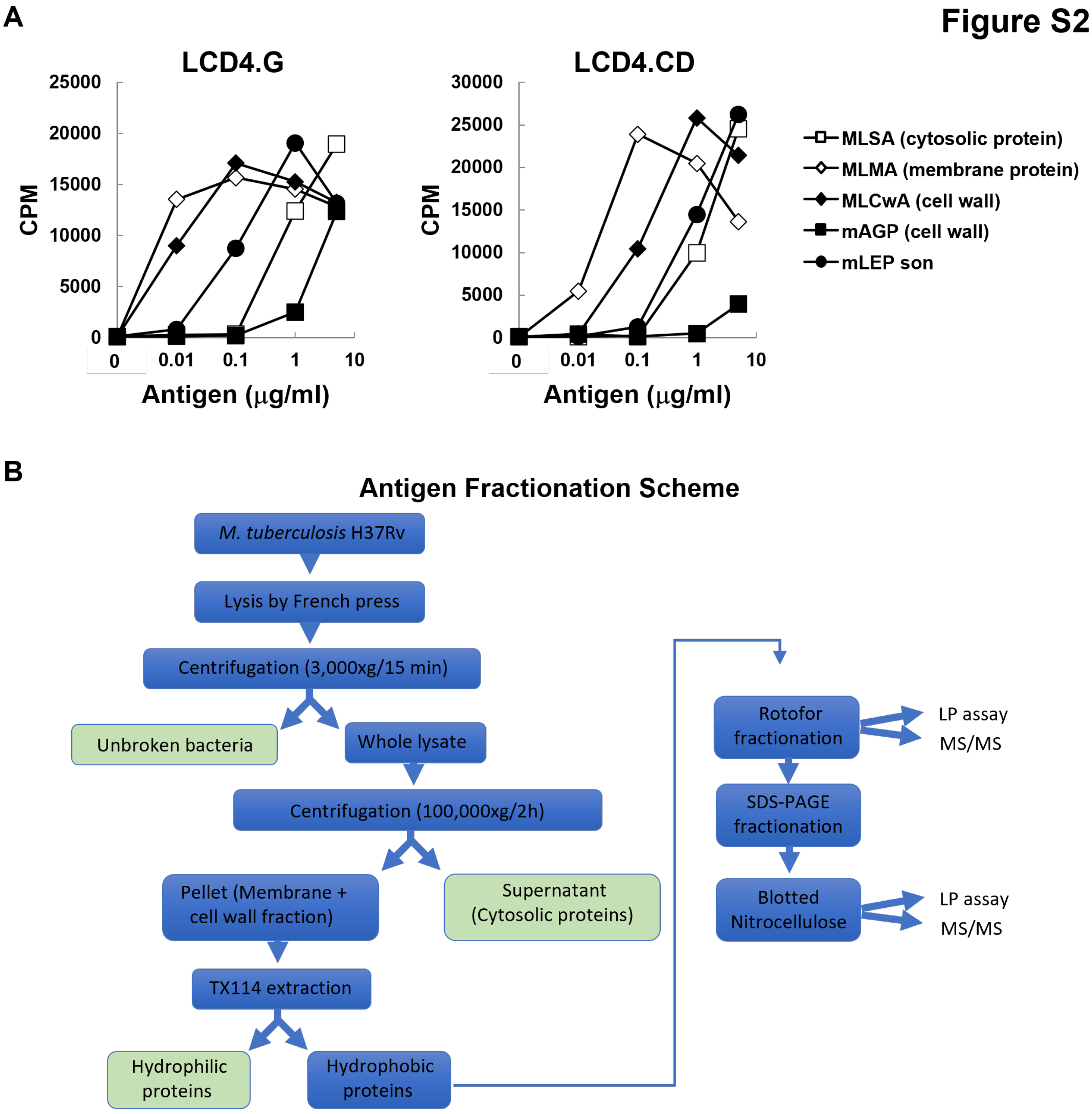
A
LCD4.G
LCD4.CD
CPM
Antigen (μg/ml)
0
0.01
0.1
1
10
MLSA (cytosolic protein)
MLMA (membrane protein)
MLCwA (cell wall)
mAGP (cell wall)
mLEP son
B
Antigen Fractionation Scheme
M. tuberculosis H37Rv
Lysis by French press
Centrifugation (3,000xg/15 min)
Unbroken bacteria
Whole lysate
Centrifugation (100,000xg/2h)
Pellet (Membrane + cell wall fraction)
Supernatant (Cytosolic proteins)
TX114 extraction
Hydrophilic proteins
Hydrophobic proteins
Rotofor fractionation
SDS-PAGE fractionation
Blotted Nitrocellulose
LP assay
MS/MS

**Figure S2. T cell activation by mycobacterial protein- and lipid-rich fractions and fractionation workflow with T cell-based "Western blot" readouts, related to Figure 2.**

(A) Proliferation response of LCD4.G and LCD4.CD T cell lines to antigenic preparations derived from *M. leprae*. MLSA (*M. leprae Soluble Antigen*), MLMA (*M. leprae* Membrane Antigen), MLCwA (*M. leprae* Cell Wall Antigen), mAGP (mycolyl-arabinogalactan-peptidoglycan complex), and mLEP son (*M. leprae* sonicated). Data indicate mean of triplicate values for each T cell line and are representative of three independent experiments.

(B) Schematic of *M. tuberculosis* Triton X-114 extraction and Rotofor preparative IEF separation. LP: lymphocyte proliferation assay; MS/MS: tandem mass spectrometry.

**Figure S3**

**A**

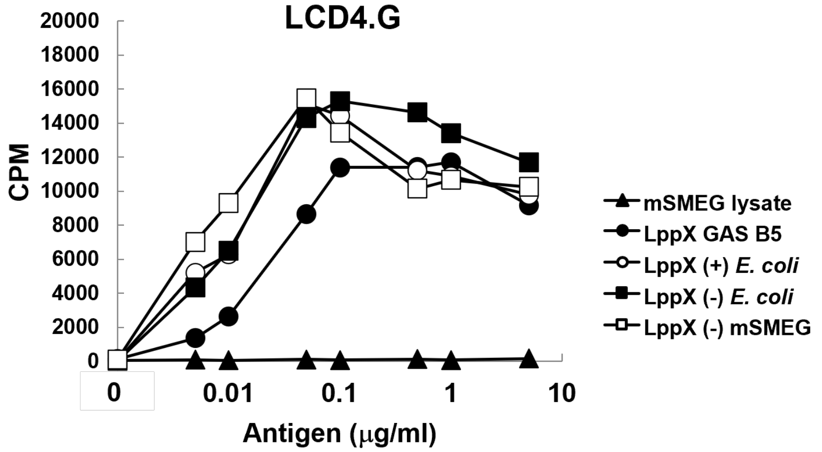


**B**

CSSPKPDAEEQGVPVSPTASDPALLAEIRQSLDATKGLTSVHVAVRTTGKVDSLLGITSADVDVRANPLAAKGVCTY
NDEQGVPFRVQGDNISVKLFDDWSNLGSISELSTSRVLDPAAGVTQLLSGVTNLQAQGTEVIDGISTTKITGTIPASSV
KMLDPGAKSARPATVWIAQDGSHHLVRASIDLGSGSIQLTQSKWNEPVNVDKLHHHHHH

Theoretical Mw: 22577.25()

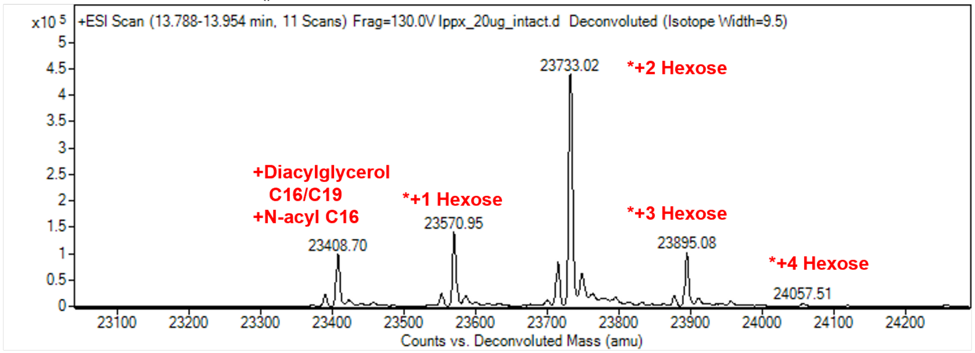


23408.70-22577.25=831.45
Modification:
+Diacylglycerol C16/C19
+N-acyl C16

**C**

MSSPKPDAEEQGVPVSPTASDPALLAEIRQSLDATKGLTSVHVAVRTTGKVDSLLGITSADVDVRANPLAAKGVCTYN
DEQGVPFRVQGDNISVKLFDDWSNLGSISELSTSRVLDPAAGVTQLLSGVTNLQAQGTEVIDGISTTKITGTIPASSVK
MLDPGAKSARPATVWIAQDGSHHLVRASIDLGSGSIQLTQSKWNEPVNVDKLAAALEHHHHHH

Theoretical Mw: 23060.62 (without signal sequence)

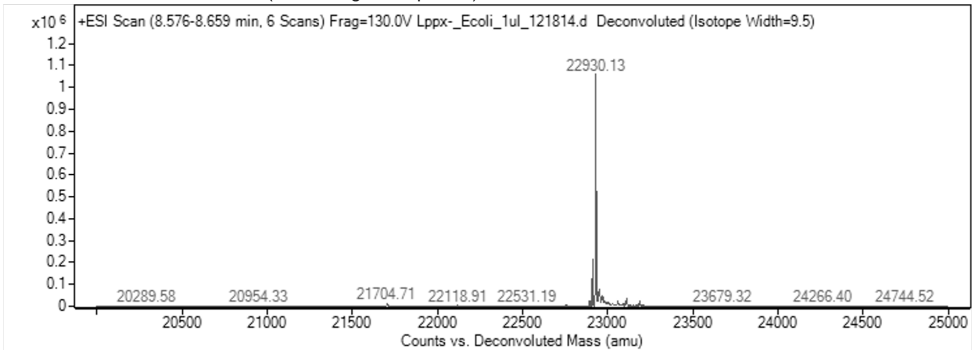

**Figure S3. T cell recognition of LppX is independent of lipidation and glycosylation and is driven by the peptide backbone.**

(A) T cell stimulatory activity of non-lipidated and/or non-glycosylated recombinant LppX expressed in *M. smegmatis* (mSMEG) or *E. coli*. Data indicate mean of triplicate values for LCD4.G T cell line and are representative of three independent experiments.

(B–C) Intact-mass and peptide-mass fingerprinting by LC–MS/MS showing (B) recombinant LppX protein (theoretical mass 22,577 Da) produced in *M. smegmatis* and (C) recombinant LppX (theoretical mass 23,060 Da) without the signal sequence (-) produced in *E. coli*. rLppX (-) lacks added acyl chains or glycans, resulting in the absence of the native 831.45 Da triacylation present in the native protein.

### Figure S4

**A**

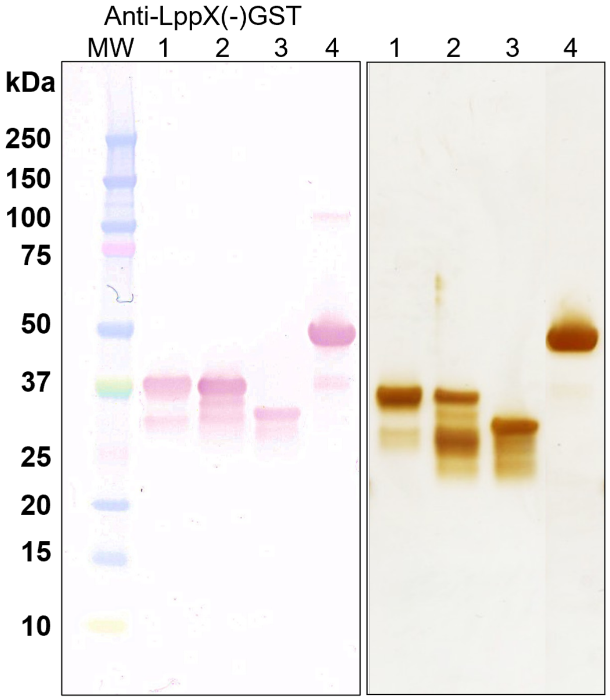


1 - LppX(-)GST-B – 35.7 kDa
2 - LppX(-)GST-C – 37.4 kDa
3 - LppX(-)GST-E – 32.6 kDa
4 - LppX(-)GST from ClearColi

**B**

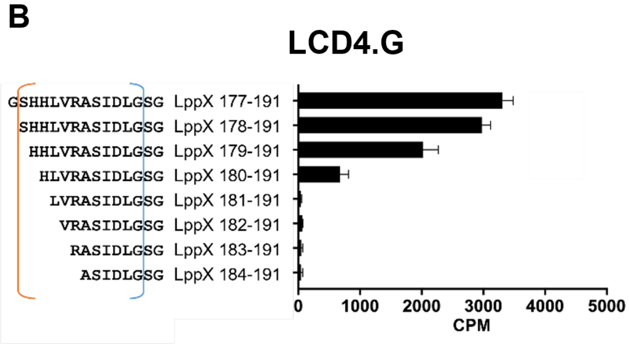

**Figure S4. C-terminal localization of the LppX T cell epitope defined by fragment mapping and truncation analysis, related to Figure 3.**

(A) Western blot analysis of recombinant LppX fragments fused to GST. Ponceau S- and Silver-stained membranes are shown.

(B) Overlapping peptides derived from the C-terminus of LppX (15–8 amino acids in length) were screened for their ability to stimulate the LCD4.G T cell line. Peptides upstream or downstream of the minimal LppX_12mer sequence (SHHLVRASIDLG) did not activate T cells. Data indicate mean ± SEM of triplicate values and are representative of three independent experiments.

### Figure S5

A

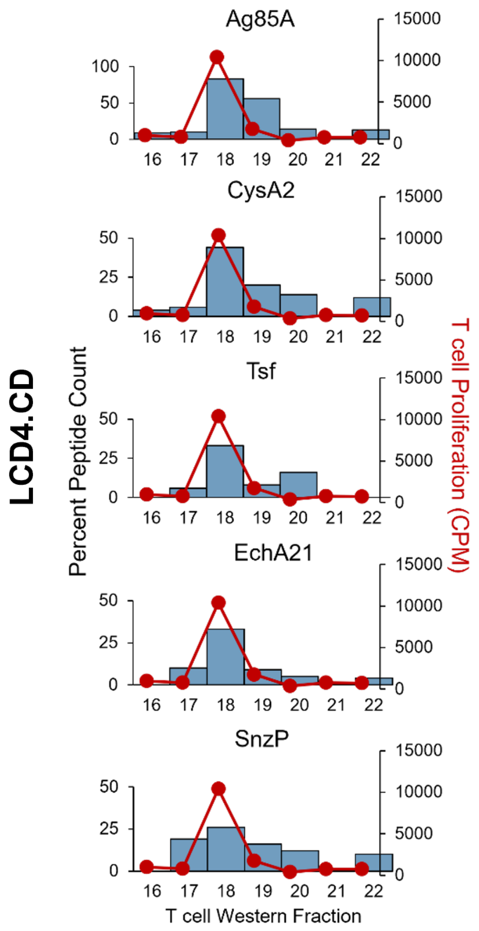


B

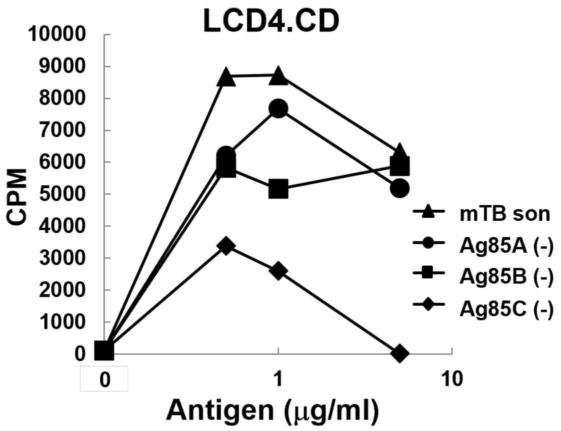


C

```
Ag85A       FSRPGLP VEYLQVPSPS MGRDIKVQFQ SGGANSPALY LLDGLRAQDD FSGWDINTPA
Ag85A-G     FSRPGLP VEYLQVPSPS MGRDIKVQFQ SGGANSPALY LLDGLRAQDD FSGWDINTPA
Ag85A-H     FSRPGLP VEYLQVPSPS MGRDIKVQFQ SGGANSPALY LLDGLRAQDD FSGWDINTPA
Ag85A-I     FSRPGLP VEYLQVPSPS MGRDIKVQFQ SGGANSPALY LLDGLRAQDD FSGWDINTPA

FEWYDQSGLS VVMPVGGQSS FYSDWYQPAC GKAGCQTYKW ETFLTSELPG WLQANRHVKP TGSAVVGLSM
FEWYDQSGLS VVMPVGGQSS FYSDWYQPAC GKAGCQTYKW ETFLTSELPG WLQANRHVKP TGSAVVGLSM
FEWYDQSGLS VVMPVGGQSS FYSDWYQPAC GKAGCQTYKW ETFLTSELPG WLQANRHVKP TGSAVVGLSM
FEWYDQSGLS VVMPVGGQSS FYSDWYQPAC GKAGCQTYKW ETFLTSELPG WLQANRHVKP TGSAVVGLSM

AASSALTLAI YHPQQFVYAG AMSGLLDPSQ AMGPTLIGLA MGDAGGYKAS DMWGPKEDPA WQRNDPLLNV
AASSALTLAI YHPQQFVYAG AMSGLLDPSQ AMGPTLIGLA MGDAGGYKAS DMWGPKEDPA WQRNDPLLNV
AASSALTLAI YHPQQFVYAG AMSGLLDPSQ AMGPTLIGLA MGDAGGYKAS DMWGPKEDPA WQRNDPLLNV
AASSALTLAI YHPQQFVYAG AMSGLLDPSQ AMGPTLIGLA MGDAGGYKAS DMWGPKEDPA WQRNDPLLNV

GKLIANNTRV WVYCGNGKPS DLGGNNLPAK FLEGFVRTSN IKFQDAYNAG GGHNGVFDFP DSGTHSWEYW
GKLIANNTRV WVYCGNGKPS DLGGNNLPAK FLEGFVRTSN IKFQDAYNAG GGHNGVFDFP DSGTHSWEYW
GKLIANNTRV WVYCGNGKPS DLGGNNLPAK FLEGFVRTSN IKFQDAYNAG GGHNGVFDFP DSGTHSWEYW
GKLIANNTRV WVYCGNGKPS DLGGNNLPAK FLEGFVRTSN IKFQDAYNAG GGHNGVFDFP DSGTHSWEYW

GAQLNAMKPD LQRALGATPN TGPAPQGA
GAQLNAMKPD LQRALGATPN TGPAPQGA
GAQLNAMKPD LQRALGATPN TGPAPQGA
GAQLNAMKPD LQRALGATPN TGPAPQGA
```

D

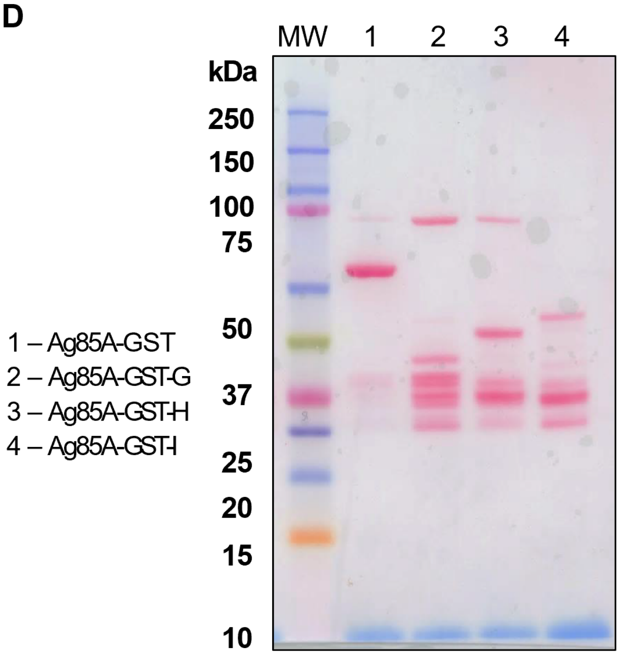

**Figure S5. Identification, validation, and cross-species conservation of an N-terminal Ag85A-derived T cell epitope, related to Figures 2 and 4.**

(A) LC-MS/MS spectra and peptide coverage maps highlighting abundant Ag85 complex peptides within LCD4.CD stimulatory IEF fractions.

(B) T cell stimulatory activity of non-lipidated and non-glycosylated recombinant Ag85A, Ag85B, and Ag85C expressed in *E. coli*. mTB son, *M. tuberculosis* sonicated. Data indicate mean of triplicate values for LCD4.CD T cell line and are representative of three independent experiments.

(C) Sequence alignment of the Ag85A protein partitioned into three segments, showing the Ag85A reference protein (blue) and the corresponding Ag85A-G (black), Ag85A-H (magenta), and Ag85A-I (green) fragments. Underlined regions indicate the boundaries of each protein segment used for GST constructs. The 12–amino acid minimal epitope is highlighted in yellow within the reference sequence.

(D) Western blot analysis of recombinant Ag85A fragments fused to GST. A Ponceau S-stained membrane is shown.

### Figure S6

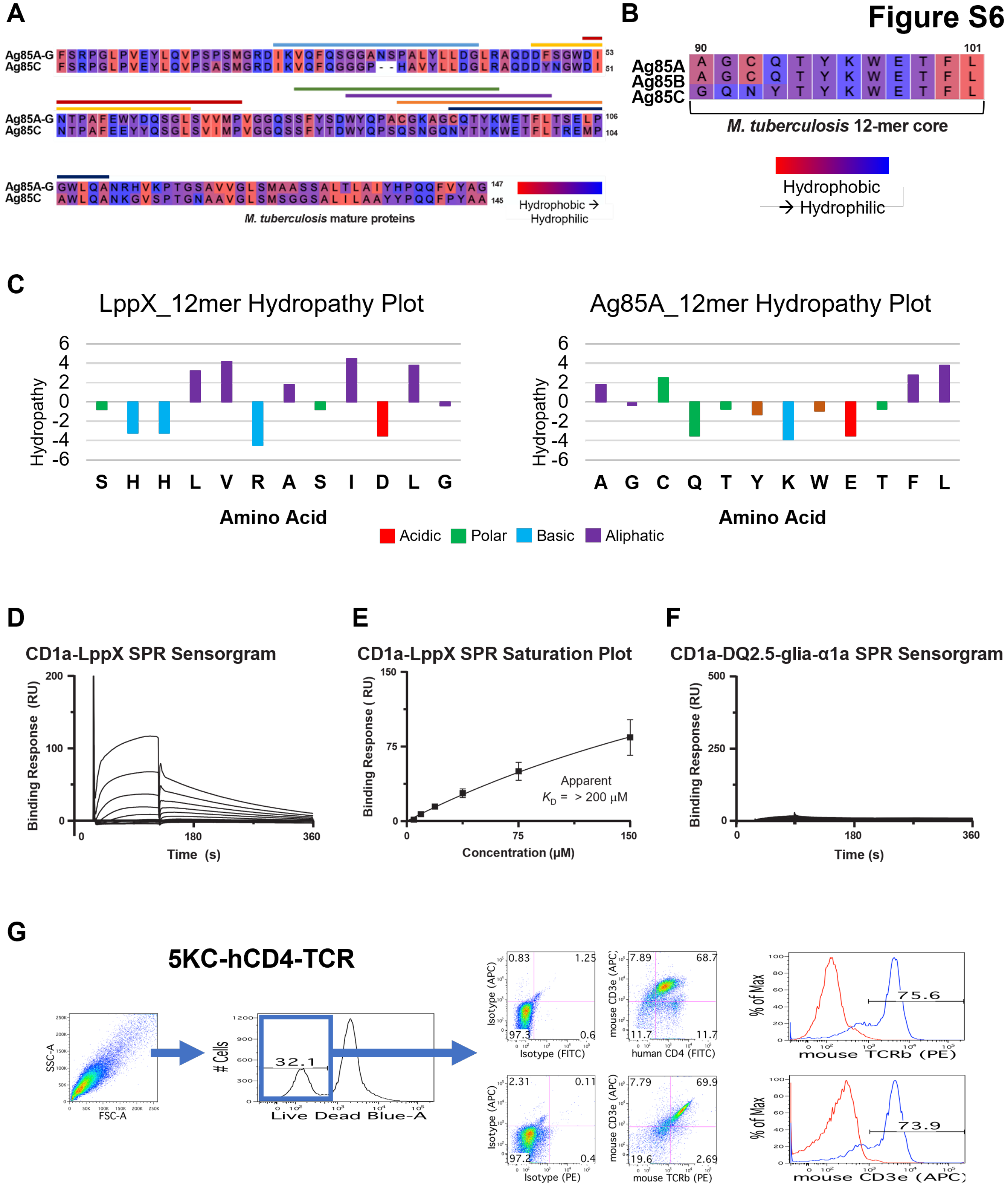

A
Ag85A-G
Ag85C
M. tuberculosis mature proteins
Hydrophobic → Hydrophilic
B
Ag85A
Ag85B
Ag85C
M. tuberculosis 12-mer core
Hydrophobic → Hydrophilic
C
LppX_12mer Hydropathy Plot
Ag85A_12mer Hydropathy Plot
Hydropathy
Amino Acid
Acidic
Polar
Basic
Aliphatic
D
CD1a-LppX SPR Sensorgram
Binding Response (RU)
Time (s)
E
CD1a-LppX SPR Saturation Plot
Binding Response ( RU)
Concentration (μM)
Apparent $K_D$ = > 200 μM
F
CD1a-DQ2.5-glia-α1a SPR Sensorgram
Binding Response (RU)
Time (s)
G
5KC-hCD4-TCR
SSC-A
FSC-A
### Cells
Live Dead Blue-A
32.1
Isotype (APC)
Isotype (FITC)
mouse CD3e (APC)
human CD4 (FITC)
Isotype (PE)
mouse TCRb (PE)
% of Max
mouse TCRb (PE)
75.6
mouse CD3e (APC)
73.9

**Figure S6. Structural and functional characterization of LppX- and Ag85A-derived CD1a-restricted T cell epitopes, related to Figures 5 and 6.**

(A) Sequence alignment of Ag85A segment G (Ag85A-G), used for GST constructs, with the corresponding region of Ag85C from *M. tuberculosis*. Underlined regions denote Ag85A-derived 20-mer peptides from the N-terminal fragment tested for activation of LCD4.CD.

(B) Alignment of the Ag85A-derived 12-mer epitope with the corresponding sequences in Ag85B and Ag85C from *M. tuberculosis*.

(C) Kyte–Doolittle hydropathy plots for LppX_12mer and Ag85A_12mer.

(D–E) Representative sensorgram and equilibrium fit (saturation plot) for CD1a binding to LppX_12mer. Error bars indicate mean ± SEM from three independent experiments. LppX_12mer showed qualitative CD1a binding at high concentrations, but a reliable $K_D$ could not be determined.

(F) Representative sensorgram for the DQ2.5-glia-α1a negative-control peptide.

(G) Flow cytometry analysis of 5KC reporter cells stably expressing human CD4 and the LCD4.G TCR. Data are representative of three independent experiments.

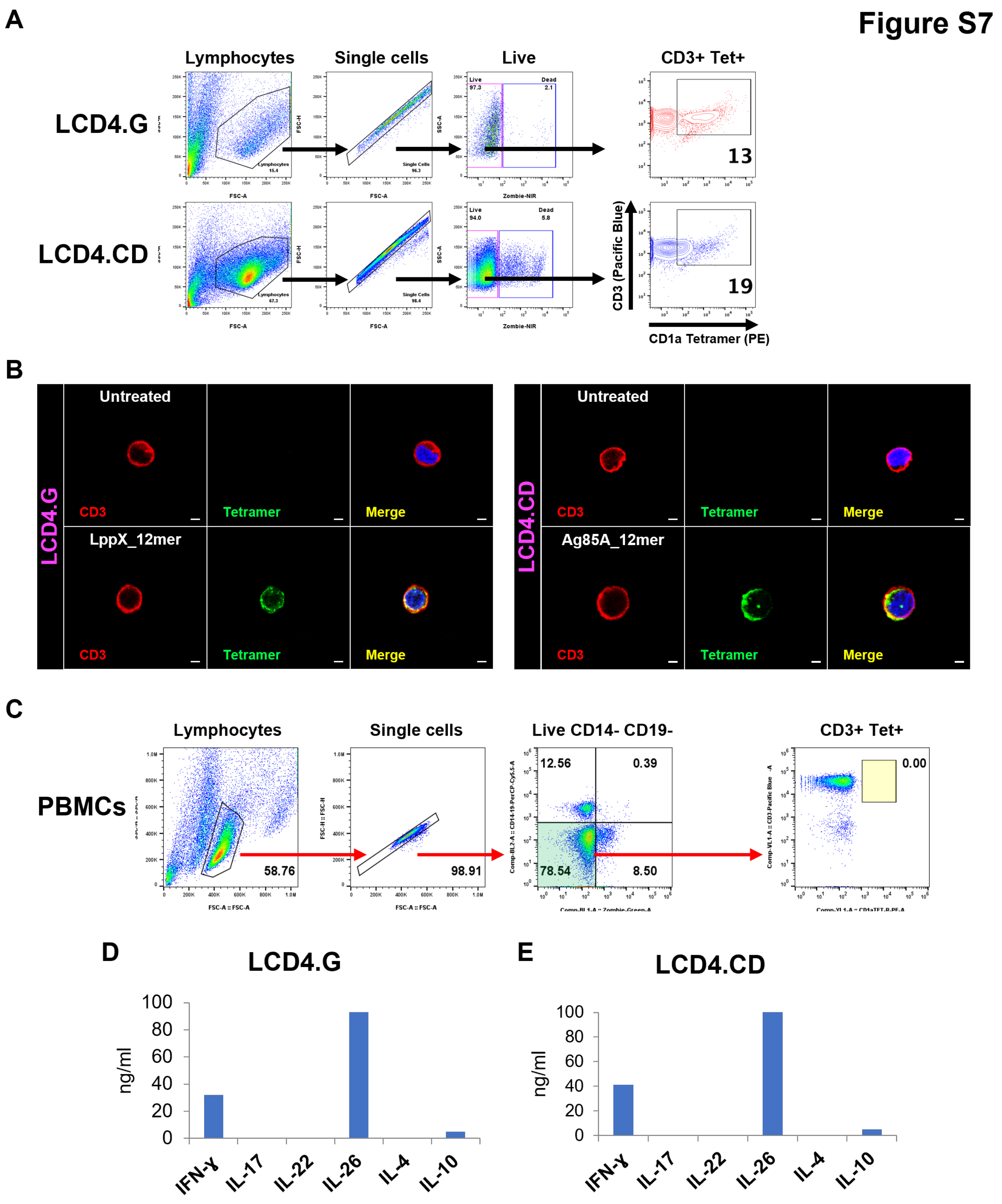
Figure S7
A
Lymphocytes
Single cells
Live
CD3+ Tet+
LCD4.G
LCD4.CD
13
19
CD3 (Pacific Blue)
CD1a Tetramer (PE)
B
Untreated
LppX_12mer
Ag85A_12mer
CD3
Tetramer
Merge
C
Lymphocytes
Single cells
Live CD14- CD19-
CD3+ Tet+
PBMCs
58.76
98.91
12.56
0.39
78.54
8.50
0.00
D
LCD4.G
E
LCD4.CD
ng/ml
IFN-γ
IL-17
IL-22
IL-26
IL-4
IL-10

**Figure S7. Identification and functional profiling of CD1a-restricted peptide-specific T cells by tetramer staining and cytokine analysis, related to Figure 7.**

(A) Flow cytometry analysis of LCD4.G and LCD4.CD T cell lines stained with Ag85A- or LppX-treated tetramers, respectively. CD1a-tetramer$^+$ cells were gated on live (Zombie negative), single, CD3$^+$ lymphocytes. Data are representative of four (LCD4.G) or five (LCD4.CD) independent experiments.

(B) Confocal microscopy analysis of LCD4.G and LCD4.CD T cell lines stained with indicated CD1a tetramers (green) treated with LppX_12mer, Ag85A_12mer, or untreated control. T cells were identified with anti-CD3 (red) mAb and nuclei (blue) were counter-stained with NucRed. Data are representative of three independent experiments for each T cell line. Scale bars = 50 μm.

(C) Flow cytometry analysis of human PBMC samples from leprosy patients (RR and L-lep) and healthy donors. CD1a-tetramer$^+$ CD3$^+$ T cells were gated on single, live (Zombie negative), CD14$^-$CD19$^-$ cells. Data are representative of eight HD, eight RR and three L-lep donors.

(D–E) Secreted cytokines produced by the LCD4.G (D) and LCD4.CD (E) T cell lines following anti-CD3/CD28 stimulation. Data indicate mean of triplicate values for each T cell line and are representative of three independent experiments.

**Table S1. Identification of LppX ortholog in *M. leprae*, related to Figure 3.**
The gene identified in Mycobrowser as LppX ortholog in *M. leprae* (MLEP) was subjected to BLAST against *M. tuberculosis* (MTB; #Rv2945c) using the model organism (landmark) database. The percentage amino acid identity and similarity (I/S) derived from BLAST analysis are summarized below, together with the number of identical amino acids within the LCD4.G peptide (LppX_12mer). The 12_mer sequences are colored in green to show the peptide homology.

| Mycobrowser | Gene # | MTB LppX (I/S) | LCD4.G Peptide | Note |
|---|---|---|---|---|
| **LppX** | MLEP ML0136 | 76/85 | 12 of 12 | Figure 3F |

```
>Mycobacterium tuberculosis H37Rv|Rv2945c|lppX

MNDGKRAVTSAVLVVLGACLALWLSGCSSPKPDAEEQGVPVSPTASDPALLAEIRQSLDATKGLTSVHVAVRTTGKV
DSLLGITSADVDVRANPLAAKGVCTYNDEQGVPFRVQGDNISVKLFDDWSNLGSISELSTSRVLDPAAGVTQLLSGV
TNLQAQGTEVIDGISTTKITGTIPASSVKMLDPGAKSARPATVWIAQDGSHHLVRASIDLGSGSIQLTQSKWNEPVN
VD

>Mycobacterium leprae TN|ML0136|lppX

MNDRKWVTSSVMLVTLSACLALGLSGCSSTKPDAQEQSSSSSPASSDPALTAEIKQSLETTKALSSVHVVVQTTGKV
DALLGISNADVDVQANPLAVKGTCTYNDQPGVPFRVLGDNISVKLFDDWSNLGSISDLSTSHVLDPNTGITQVLSGV
INLQAQGTEVVDRIPTNKITGTVPTSSVKMLDPKAKGSKLATVWIAQDGSHHLVRASIDLGSGSIQLTQSKWNEPVN
TN
```

**Table S2. Identification of Ag85A homologs in *M. smegmatis*, related to Figure 4.**
The genes identified in Mycobrowser as Ag85 homologs in *M. smegmatis* (MSMEG) were subjected to BLAST against *M. tuberculosis* (MTB; #004032) using the model organism (landmark) database. The percentage amino acid identity and similarity (I/S) derived from BLAST analysis are summarized below, together with the number of identical amino acids within the LCD4.CD peptide (Ag85A_12mer) and the 10-amino acid shared sequence (CQTYKWETFL). The sequences are colored to show the peptide homology: 12_mer sequences are highlighted in green; 10-aa sequences are bolded; Amino acid changes within 10-aa are colored in red; and amino acid changes within 12_mer sequence are highlighted in yellow.

| Mycobrowser | Gene # | MTB Ag85A (I/S) | MTB Ag85B (I/S) | MTB Ag85C (I/S) | LCD4.CD 12mer Peptide | LCD4.CD 10-aa Peptide | Note |
|---|---|---|---|---|---|---|---|
| **Antigen 85-A (fbpA)** | MLEP ML0097 | 83/89 | 77/86 | 65/75 | 12 of 12 | 10 of 10 | Figure 4F |
| **Antigen 85-A** | MSMEG 6398 | 69/79 | 68/80 | 64/77 | 10 of 12 | 9 of 10 | UniProt/TR data |
| **Antigen 85-B** | None | N/A | N/A | N/A | N/A | N/A | |
| **Antigen 85-C** | MSMEG 2078 | 69/81 | 71/81 | 70/80 | 10 of 12 | 9 of 10 | UniProt/SP data; Figure 4F |
| **Antigen 85-C** | MSMEG 3580 | 66/78 | 68/77 | 75/84 | 8 of 12 | 8 of 10 | |
| **Antigen 85-C** | MSMEG 6396 | 41/55 | 39/53 | 41/52 | 5 of 12 | 5 of 10 | Best Homology to MPT51 (67/78) |
| **Antigen 85-C** | MSMEG 6399 | 55/67 | 57/68 | 59/70 | 8 of 12 | 8 of 10 | |
| **Antigen 85-C** | MSMEG 6583 | 64/76 | 64/76 | 67/78 | 10 of 12 | 8 of 10 | |

>Mycobacterium tuberculosis MTBC0|mtbc0_004032|ag85A

MQLVDRVRGAVTGMSRRLVVGAVGAALVSGLVGAVGGTATAGAFSRPGLPVEYLQVPSPSMGRDIKVQFQSGGANSP
ALYLLDGLRAQDDFSGWDINTPAFEWYDQSGLSVVMPVGGQSSFYSDWYQPACGKAG**CQTYKWETFL**TSELPGWLQA
NRHVKPTGSAVVGLSMAASSALTLAIYHPQQFVYAGAMSGLLDPSQAMGPTLIGLAMGDAGGYKASDMWGPKEDPAW
QRNDPLLNVGKLIANNTRVWVYCGNGKPSDLGGNNLPAKFLEGFVRTSNIKFQDAYNAGGGHNGVFDFPDSGTHSWE
YWGAQLNAMKPDLQRALGATPNTGPAPQGA

>Mycobacterium leprae TN|ML0097|fbpA

MKFVDRFRGAVAGMLRRLVVEAMGVALLSALIGVVGSAPAEAFSRPGLPVEYLQVPSPSMGRDIKVQFQNGGANSPA
LYLLDGLRAQDDFSGWDINTTAFEWYYQSGISVVMPVGGQSSFYSDWYSPACGKAG**CQTYKWETFL**TSELPQYLQSN
KQIKPTGSAAVGLSMAGLSALTLAIYHPDQFIYVGSMSGLLDPSNAMGPSLIGLAMGDAGGYKAADMWGPSTDPAWK
RNDPTVNVGTLIANNTRIWMYCGNGKPTELGGNNLPAKLLEGLVRTSNIKFQDGYNAGGGHNAVFNFPDSGTHSWEY
WGEQLNDMKPDLQQYLGATPGA

>Mycobacterium smegmatis MC2–155|MSMEG_6398|MSMEG_6398

MKFVGRMRGAAAGLSRRLTVAVAAAAVLPGLVGVVGGSATAGAWSRPGLPVEYLEVPSAAMGRDIRVEFQSGGPGAP
ALYLLDGMRAREDQNGWDIELPTFEWFLNSGISVVMPVGGQSSFYSDWYKPACGSKDGG**CKTYKWETFL**TQELPAWL

AANRDVKPTGSAVVGLSMAGSSALMLAARHPQQFIYAASLSGTLNPSEGWWPMLIGISMGDAGGYKADDMWGSTNDP
NNAWKANDPTENVATIANNGTRIWVYCGNGKPGELGGTDLPAKFLEGFVCRTNSTFQEKYIEAGGKNGVFNFPQSGT
HNWAYWGQQLQAMKPDLQRVLGATPTA

>Mycobacterium smegmatis MC2–155|MSMEG_2078|MSMEG_2078

MTFIDKIRGHWARRMTVAAVAALLLPGLVGVVGGSATAGAFSRPGLPVEYLMVPSPSMGRDIKVQFQSGGPGSHAVY
LLDGLRAQDDFNGWDINTNAFEMFLDSGLSVVMPVGGQSSFYSDWYQPACGNN**GCVTYKWETFL**TSELPEWLAANRD
VAATGNAAIGLSMAGSAALILAAYHPDRFIYAGSMSGFLNPSEGWWPFLINISMGDAGGYKANDMWGPTEDPNSAWK
RNDPMVQIPRLVANNTRIWVYCGNGQPNELGGGDLPATFLEGLTIRTNETFRDNYIAAGGNNGVFNFPNNGTHNWAY
WGRELQAMVPDLQRVLG

>Mycobacterium smegmatis MC2–155|MSMEG_3580|MSMEG_3580

MRGIAAWKALRRFVIGALAALMLPGLIGFAGGSASAGAFSRPGLPVEYLDVFSPSMNRDIRVQFQGGGPHAVYLLDG
LRAQDDYNGWDINTPAFEWFYQSGLSTIMPVGGQSSFYTDWYQPSKGNGQ**DYTYKWETFL**TQELPAWLEANRGVSRT
GNAVVGLSMAGSAALTYAIYHPEQFIYASTLSGFLNPSEGWWPMLIGLAMNDAGGYNAESMWGPSTDPAWKRNDPMV
NINQLVANNTRLWVYCGTGTPSELDAGVSGGNLLAAQFLEGLTLRTNLTFRDQYIAAGGTNAVFNFPPNGTHTWNYW
GQQLLEMKPDIQRVLGAQSAT

>Mycobacterium smegmatis MC2–155|MSMEG_6396|MSMEG_6396

MRRGLSLVRALMLTVVLAAGLWTVSATSGAPARADGVEYLMVPSAAMGRDIPVAFQAGGPHAVFLLDAFNAAPDVSN
WVNAGSAMSTLAGRGISVAAPAGGAWSLYTNWEQ**DGSKQWETFL**TAELPGWLAANKGLAPDGHAVVGAAQGGTAAVT
LAAFHPGMFRFAGSLSGFLTPSATTLNGAITAGLARFGNVDANRMWGPPQFGRWKWHDPAVHVQLLADSNTRLWVYS
PGTLTCSDPAAMIGYCDQAQGSNRQFYSDYRRVGGSNGHFDFPASGQHDWGSWAPQLAYMSGELVATIK

>Mycobacterium smegmatis MC2–155|MSMEG_6399|MSMEG_6399

MAPMVSTRAVRVSGLRLLVATVISVMFTTFAVASLEIPRANAFSREGLPVEYLDVYSPSMGRNLRVQFQGPGETVDD
MRPSRAVYLLDGLRAQDDYSGWDINTPAFEWFYNSGISVVMPVGGQSSFYSDWYSPSSFNNQ**TYTYKWETFL**TRELP
AWLAANRNVSTTGNGVVGLSMSGSAALILSAFHPGQFRYAASLSGFLNPSSLLMQQAIRVAMLDAGGYNVDNMWGPP
WDGAWKRNDPIRQVDRIVANGTRLWIYCAPGGATPLDDGADPNLAMSANSLETLAIKSNKDFQEAYVRAGGRNATFT
FPPAGNHAWPYWGAQLSALKPDLIASLNG

>Mycobacterium smegmatis MC2–155|MSMEG_6583|MSMEG_6583

MKPSFFSTLSRRVAAGAATVLMLAGLIATGPVPQAAAYSRDGLPVERLQVPSAAMGRDITVQFQGGGPHALYLLDGL
RAQDDANGWDINTAAFEWFYQSGISVVMPVGGQSSFYTDWYRPAVGSAG**TTTYKWETFL**TRELPAWLAANRGVEPTG
NAVVGLSMSGGAALNLATWYPQQFIFAGALSGFLNPSQGLWPTMIGFAMKDAGGYNSADMWGLANDPAWRRNDPMVN
INRLVANNTAIWVYCGNGAPSDLDAAGDFGQLYSAQFLENITVNTNKEFQKRYLAAGGHNAVFNFPPNGTHAWGYWG
AQLQAMKPDLLRVLGVGAPAPAAPAAPAPAAPVPAAPGMPVAPVAPAVPGAQGLPAAPAYPVAPGVQGVPGVQGVPA
APASPAVALPATRPV

**Table S3. Clinical characteristics of L-lep and RR patient groups, related to Figure 7.**

| | Specimen ID | Clinical Form | Gender | Nationality | Ethnicity | Age | BI | SBI |
|---|---|---|---|---|---|---|---|---|
| L-Lep Group | **LL1** | LL | M | Brazilian | Mixed | 16 | 5 | 5.8 |
| | **LL2** | LL | M | Brazilian | Mixed | 38 | 4 | 5.9 |
| | **LL3** | LL | F | Brazilian | Black | 25 | 5.25 | 5.8 |
| RR Group | **RR1** | BL | M | Mexican | Hispanic | 49 | N/A | N/A |
| | **RR2** | BL | M | Filipino | Asian | 49 | N/A | N/A |
| | **RR3** | LL | M | Mexican | Hispanic | 78 | N/A | N/A |
| | **RR4** | BL | F | Filipino | Asian | 35 | N/A | N/A |
| | **RR5** | BL | M | Filipino | Asian | 65 | N/A | N/A |
| | **RR6** | LL | M | Filipino | Asian | 59 | N/A | N/A |
| | **RR7** | LL | M | American | White | 66 | N/A | N/A |
| | **RR8** | BL | M | Chinese | Asian | 65 | N/A | N/A |

BL= Borderline-Lepromatous; LL= Lepromatous-Lepromatous; RR = Reversal Reaction; BI = Bacillary Index; SBI = Skin Bacillary Index; N/A = not applicable
